# Thymidine rescues ATR kinase inhibition induced deoxyuridine contamination in genomic DNA, cell death, and Type 1 interferon expression

**DOI:** 10.1101/2022.02.24.481821

**Authors:** Norie Sugitani, Frank P. Vendetti, Andrew J. Cipriano, Pinakin Pandya, Joshua J. Deppas, Tatiana N. Moiseeva, Sandra Schamus-Haynes, Yiyang Wang, Drake Palmer, Hatice U. Osmanbeyoglu, Anna Bostwick, Nathaniel W. Snyder, Yi-Nan Gong, Katherine M. Aird, Greg M. Delgoffe, Jan H. Beumer, Christopher J. Bakkenist

## Abstract

ATR kinase is a central regulator of the DNA damage response (DDR) and cell cycle checkpoints. ATR kinase inhibitors (ATRi’s) combine with radiation to generate CD8^+^ T cell-dependent responses in mouse models of cancer. We show that ATRi’s induce CDK1-dependent origin firing across active replicons in CD8^+^ T cells activated *ex vivo* while simultaneously decreasing the activity of rate-limiting enzymes for nucleotide biosynthesis. These pleiotropic effects of ATRi induce deoxyuridine (dU) contamination in genomic DNA, R loops, RNA-DNA polymerase collisions, and type-1 interferons (IFN-1). Remarkably, thymidine rescues ATRi-induced dU contamination, cell death, and IFN-1 expression in proliferating CD8^+^ T cells. Thymidine also rescues ATRi-induced cancer cell death. We propose that ATRi-induced dU contamination contributes to dose-limiting leukocytopenia and inflammation in the clinic and CD8^+^ T cell dependent anti-tumor responses in mouse models. We conclude that ATR is essential to limit dU contamination in genomic DNA and IFN-1 expression.

## INTRODUCTION

The DNA Damage Response (DDR) is a signaling system that integrates DNA repair with the cell cycle to safeguard genome stability (Ciccia and Elledge, 2010). ATR is an essential DDR kinase activated at single-stranded DNA associated with stalled and collapsed DNA replication forks and resected DNA double strand breaks (Brown and Baltimore, 2000; Cortez et al., 2001; Zou and Elledge, 2003). ATR phosphorylates and activates CHK1 kinase which phosphorylates and inactivates CDC25A phosphatase (Liu et al., 2000; Mailand et al., 2000). Since CDC25A dephosphorylates and activates Cyclin-dependent kinases (CDKs) that drive the cell cycle, loss of CDC25A activity causes cell cycle arrest (Hoffmann et al., 1994). While DNA damage-induced ATR-CHK1-CDC25A signaling to cell cycle checkpoints is well-accepted, the significance of this signaling in unperturbed S phase cells is less clear (Sorensen et al., 2004).

ATR kinase inhibitors have advanced to phase 1 and phase 2 trials: ceralasertib (AZD6738); berzosertib (M6620, VX-970); elimusertib (BAY 1895344); and RP-3500 (we use ATRi’s for these four inhibitors) (Foote et al., 2018; Hall et al., 2014; Roulston et al., 2021; Wengner et al., 2020). ATR is an attractive target as cancer cells with inactivating mutations in ARID1A, ATM, ERCC1, RNASE H2, and XRCC1 are sensitive to ATRi’s *in vitro* and *in vivo*, particularly in combination with DNA damaging agents that target DNA replication forks (Hustedt et al., 2019; Mohni et al., 2014; Mohni et al., 2015; Vendetti et al., 2015; Wang et al., 2019; Williamson et al., 2016). Furthermore, ATR inhibitor AZD6738 (ATRi) combines with radiation to generate CD8^+^ T cell-dependent responses in immunoproficient mouse models of cancer (Dillon et al., 2019; Sheng et al., 2020; Vendetti et al., 2018). In the clinic, ATRi suppresses circulating monocytes and proliferating T cells and these populations rebound when ATRi clears (Krebs et al., 2018). These data suggest that the impact of ATRi’s in immune cells may be clinically important.

ATRi’s have identified ATR kinase signaling in unperturbed cells that is below the detection of phosphospecific antibodies. In one line of investigation, ATR kinase inhibition was found to induce CDK1-dependent phosphorylation on RIF1 serine-2205 and origin firing across active replicons (Moiseeva et al., 2017; Moiseeva et al., 2019b). In a second line of investigation, Françoise Bontemps and Caius Radu found that ATR kinase inhibition reduces the activities of deoxycytidine kinase (dCK) and ribonucleotide reductase (RNR), respectively, rate-limiting enzymes for nucleotide biosynthesis (Amsailale et al., 2012; Le et al., 2017). While ATR kinase inhibition decreased an ATR phosphorylation on dCK serine-74 that is associated with dCK activity, ATR kinase inhibition increased a CDK1/CDK2 phosphorylation on RRM2 threonine-33 and decreased the amount of this RNR subunit (Amsailale *et al*., 2012; D’Angiolella et al., 2012; Koppenhafer et al., 2020; Le *et al*., 2017).

We investigated the impact of ATRi on CD8^+^ T cells activated *ex vivo,* a clinically relevant model of non-immortalized cell proliferation. We show that ATRi induces deoxyuridine (dU) contamination in genomic DNA, R loops, RNA-DNA polymerase collisions, IFN-1 expression, and death in proliferating CD8^+^ T cells. We show that thymidine (dT) rescues ATRi-induced dU contamination, cell death, and IFN-1 expression in proliferating CD8^+^ T cells. We also show that dT rescues ATRi-induced cancer cell death, indicating that the underlying mechanisms are not unique to proliferating CD8+ T cells. Our data reveal that ATR kinase signaling integrates nucleic acid metabolism with the cell cycle to safeguard genome stability.

## RESULTS

### ATR kinase activity is essential in proliferating CD8^+^ T cells

We investigated the impact of ATRi on the activation, proliferation, and survival of Pmel-1 CD8^+^ T cells activated with the peptide gp100 in splenocytes when the endpoint allowed the identification of CD8^+^ T cells. We used CD8^+^ T cells purified by negative selection and activated with CD3_Ɛ_ and CD28 antibodies when contaminating cells could compromise the endpoint. CD8^+^ T cells were fully activated (CD44^hi^) at ∼21 h and started dividing at ∼24 h (Figure 1A). Once activated, CD8^+^ T cells divided at least 4 times from 24-48 h, as reported previously (Yoon et al., 2010). We observed a remarkable increase in ATR, CHK1, RNR subunits RRM1 and RRM2, dUTP nucleotidohydrolase (DUT), and WEE1 protein in proliferating CD8^+^ T cells, as expected (Figure 1B and S1).

**Figure 1:**
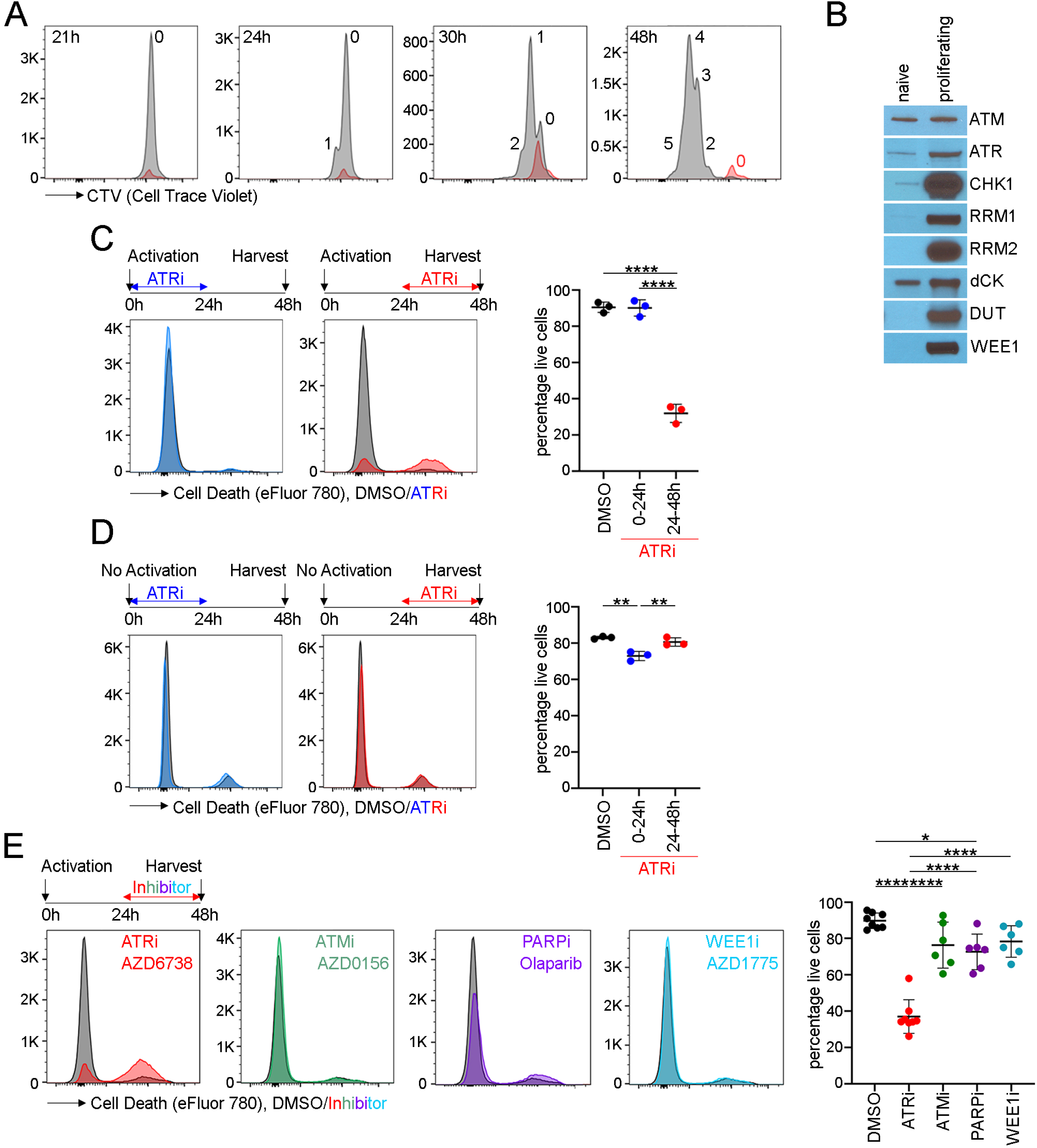
ATRi induces death in activated and proliferating CD8^+^ T cells *ex vivo*. (A) Proliferation of Pmel-1 CD8^+^ T cells activated with gp100 *ex vivo*. CTV histograms of live, CD44^hi^ CD8^+^ T cells at indicated times post-activation are shown. Red histograms are un-activated CD8^+^ T cells which have largely died. The number of divisions is shown. (B) Immunoblots of CD8^+^ T whole cell extracts prepared the same number of cells at 0 and 30 h post-activation. (C) CD8^+^ T cells were activated and treated with 5 µM AZD6738 (ATRi) from 0-24 h (blue) or 24-48 h (red). At 48 h, live CD8^+^ T cells (eFluor 780^-^CD8^+^TCRβ^+^) was quantitated. (D) Unactivated CD8^+^ T cells were maintained in IL-7 and treated with ATRi from 0-24 h or 24-48 h. At 48 h, live CD8^+^ T cells was quantitated. (E) CD8^+^ T cells were activated and treated with ATRi, 500 nM AZD0156 (ATMi), 5 μM Olaparib (PARPi), or 500 nM AZD1775 (WEE1i) from 24-48 h. At 48 h, live CD8^+^ T cells was quantitated. (C-E) Mean and SD bars shown.*:P< 0.05;**:P<0.01;****:P <0.0001 by ANOVA with Tukey’s multiple comparisons.

To determine whether ATR kinase activity is required for the activation and/or proliferation of CD8^+^ T cells, we treated CD8^+^ T cells with ATRi from 0-24 h (during activation) or 24-48 h (during proliferation) and quantitated survival using the viability dye eFluor 780. While ATRi had negligible impact on CD8^+^ T cells from 0-24 h, ATRi from 24-48 h induced death in proliferating CD8^+^ T cells (Figure 1C). We then examined whether ATRi affects the survival of un-activated, naïve CD8^+^ T cells maintained in interleukin 7 (IL-7). While ATRi caused a slight decrease in the survival of naïve CD8^+^ T cells from 0-24 h, ATRi from 24-48 h had no impact (Figure 1D). IL-7 can promote homeostatic proliferation and this may explain why ATRi caused a slight decrease in the survival of CD8^+^ T cells from 0-24 h (Tan et al., 2001). Finally, we determined whether widely used doses of clinical inhibitors of ATM (50-500 nM AZD0156), PARP (5 µM Olaparib), and WEE1 (500 nM AZD1775) induced death in proliferating CD8^+^ T cells (Moiseeva et al., 2019a; Riches et al., 2020). This is of interest because combinations of ATRi’s with these inhibitors are in clinical trials. Inhibition of PARP, but not ATM and WEE1, induced death in proliferating CD8^+^ T cells in 24 h, but this was less than the ATRi-induced CD8^+^ T cell death (Figure 1E).

### ATR kinase activity is essential in previously activated, proliferating CD8^+^ T cells *in vivo*

ATRi potentiates CD8^+^ T cell-dependent anti-tumor activity following conformal radiation in syngeneic CT26 tumors, despite transiently reducing activated CD8^+^ T cells in BALB/c host spleens (Vendetti *et al*., 2018). To define the impact of ATRi on CD8^+^ T cell activation and proliferation *in vivo*, we treated CT26 tumor-bearing mice with 75 mg/kg of ATRi on days 1,2, and 3, and immunoprofiled CD8^+^ T cells in tumor infiltrating lymphocytes (TIL) and the periphery (spleen and draining lymph node (DLN)) on day 4. ATRi reduced CD8^+^ T cell numbers in TIL >3-fold (Figure 2A). ATRi did not change relative CD8^+^ T cell numbers in the periphery. However, ATRi treatment significantly reduced spleen weight (Figure S2A), indicating that ATRi also reduced other splenic immune populations. Importantly, ATRi treatment significantly reduced the percentage of proliferating CD8^+^ T cells (Ki67^+^) in TIL, spleen, and DLN by ∼1.8-2.1-fold (Figure 2B and 2C). Therefore, ATRi impacts proliferating CD8^+^ T cells *in vivo* in all immunoprofiled tissues.

**Figure 2:**
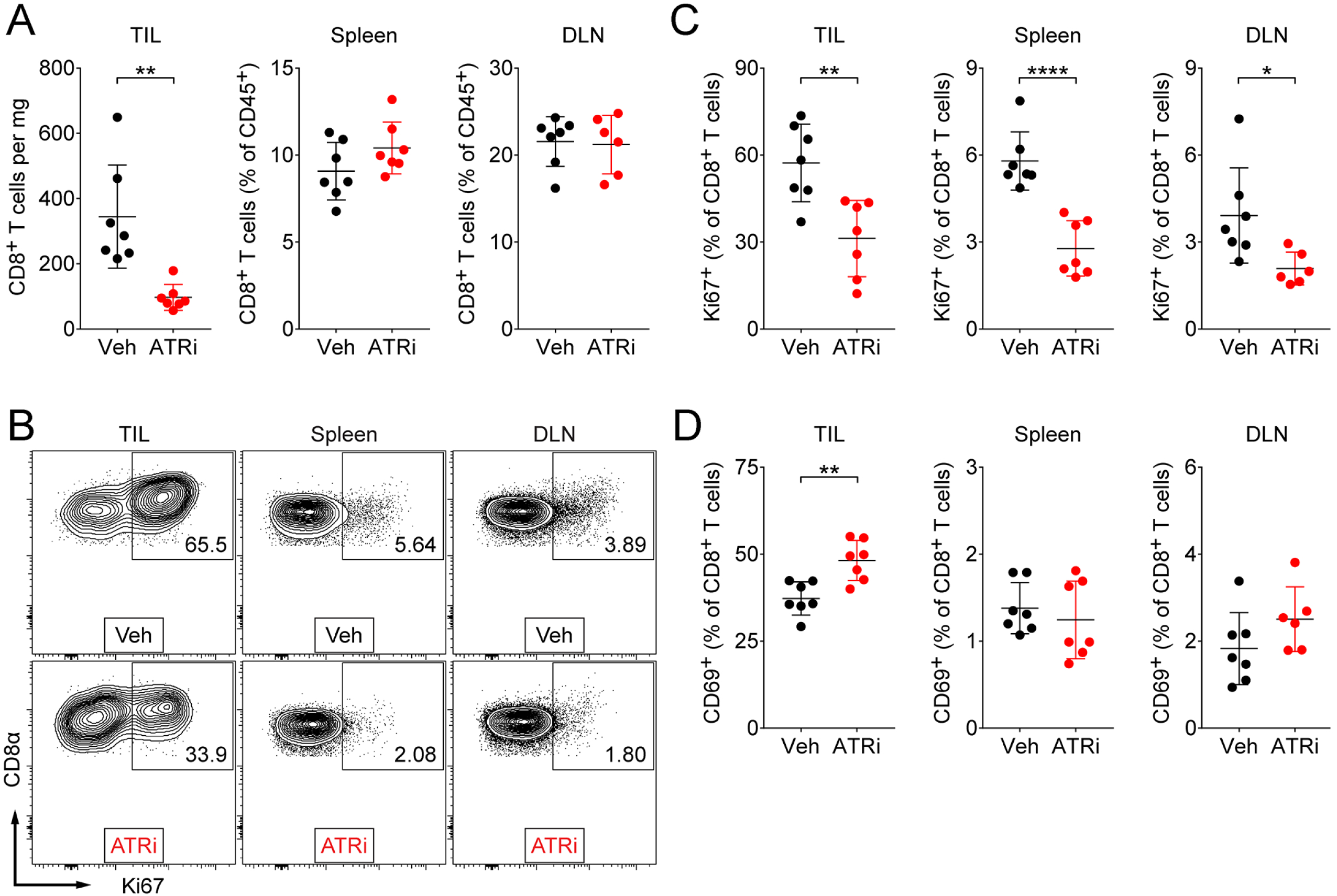
ATRi induces death in proliferating CD8^+^ T cells *in vivo.* (A-H) Tissues were harvested on day 4 from CT26 tumor-bearing mice treated with 75 mg/kg AZD6738 (ATRi) or vehicle on days 1-3 and immunoprofiled. (A) Quantitation of tumor-infiltrating (TIL) CD8^+^ T cells (per mg of tumor) and of spleen and tumor-draining lymph node (DLN) CD8^+^ T cells (percentage of total CD45^+^ cells). (B) Representative contour plots of Ki67^+^ expression in CD8^+^ T cells in the TIL, spleen, and DLN. (C) Quantitation of proliferating (Ki67^+^) CD8^+^ T cells in the TIL, spleen, and DLN. (D) Quantitation of CD69^+^ CD8^+^ T cells in the TIL, spleen, and DLN. (A-D) n=7 mice total (6 DLN for ATRi) from 2 independent experiments, each with 3-4 mice per arm. (A, C-D) Mean and SD bars shown.*:P< 0.05,**:P< 0.01,****:P< 0.0001 by two-tailed, unpaired t-test.

To determine whether ATRi impedes CD8^+^ T cell activation *in vivo*, we examined expression of the early activation marker CD69 on CD8^+^ T cells in the TIL and periphery. ATRi increased the percentage of newly or recently activated CD8^+^ T cells (CD69^+^) in TIL and this was unchanged in the periphery (Figure 2D). We further probed CD8^+^ T cell activation phenotypes using the markers CD62L and CD44 to identify un-activated naïve (TN, CD26L^lo^ CD44^lo^) CD8^+^ T cells and activated central memory (TCM, CD26L^hi^ CD44^hi^) and effector/effector memory (TEM, CD26L^lo^ CD44^hi^) CD8^+^ T cells. Consistent with our previous findings (Vendetti et al., 2018), ATRi reduced the percentage of activated TCM and TEM CD8^+^ T cells in the spleen (Figure S2B and C). Concurrently, ATRi increased the percentage of un-activated TN CD8^+^ T cells in the spleen. There were no significant changes in these populations in the DLN or TIL (Figure S2B and 2D).

### The majority of proliferating CD8^+^ T cells are in S phase

To determine the cell cycle distribution of proliferating CD8^+^ T cells, and exponentially dividing primary fibroblasts and B16 cancer cells, we labelled S phase cells with 5-ethynyl-2’-deoxyuridine (EdU) for 30 min and identified EdU incorporation and DNA content using flow cytometry. In proliferating CD8^+^ T cells, G1 was ∼1.0±0.3 h, S phase was ∼4.7±0.3 h, and G2/M was ∼0.3±0.1 h (Figure 3A and S3A-C). In fibroblasts, G1 was ∼13±3.3 h, S phase was ∼8.2 h±1.8 h, and G2/M was ∼7.5±2.3 h. Cartoons summarize the cell cycle distribution with the circumference of the circle represents the doubling time (Figure 3B and S3D). In contrast to other cell types examined, the vast majority of proliferating CD8^+^ T cells are in S phase.

**Figure 3:**
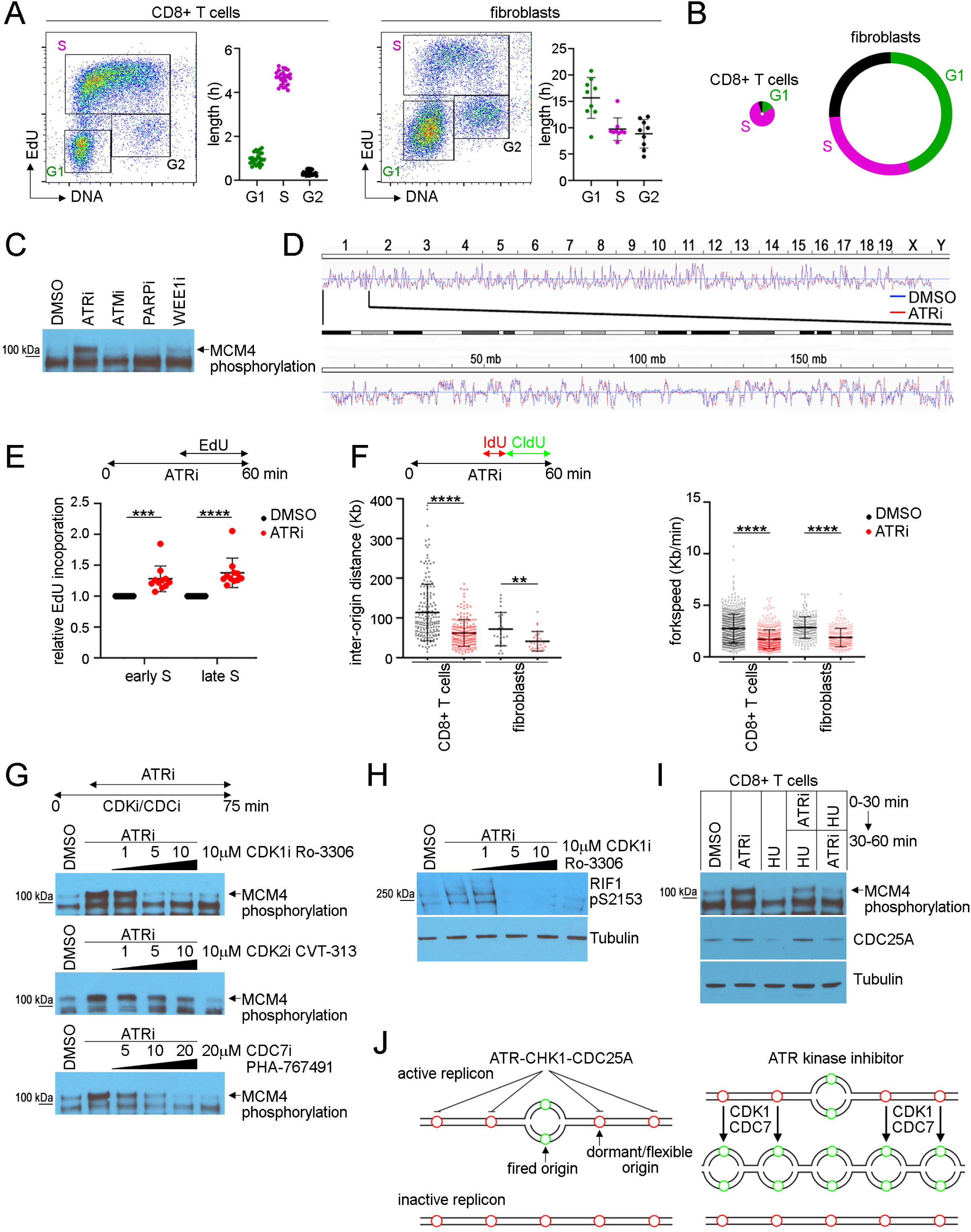
ATR kinase limits origin firing across active replicons in proliferating CD8^+^ T cells. (A) EdU versus DNA plots of CD8^+^ T cells and fibroblasts. The percentage of cells in each cell cycle phase and the doubling time was used to estimate the length of G1, S, and G2/M. (B) The circumference of the circle represents the doubling time and the length of G1, S, and G2/M is drawn to scale. (C) Immunoblot of MCM4 in the chromatin fraction prepared from CD8^+^ T cells treated with 5 μM AZD6738 (ATRi), 500 nM AZD0156 (ATMi), 5 μM Olaparib (PARPi), or 500 nM AZD1775 (WEE1i) for 1 h. (D) Repli-seq analyses of the replication timing program of CD8^+^ T cells treated with ATRi (red) or DMSO (blue) for 1 h. Upper - whole genome. Lower - chromosome 1. Data deposited at GEO (code: GSE183412). (E) EdU incorporation in early (2-3N) and late (3-4N) S phase in CD8^+^ T cells treated with ATRi for 1h. 10 µM EdU was added from 30-60 min. (F) DNA combing analyses of CD8^+^ T cells and fibroblasts treated with ATRi for 1 h. Cells were treated with IdU from 30-40 min and CldU from 40-60 min of the treatment. (G) Immunoblots of MCM4 in the chromatin fraction prepared from CD8^+^ T cells treated with increasing concentrations of Ro-3306 (CDK1i), CVT-313 (CDK2i), or PHA-767491 (CDC7i) for 75 min. Cells were treated with ATRi from 15-75 min. (H) Immunoblot of RIF1 phosphoserine-2153 in whole cell extracts prepared from CD8^+^ T cells treated with Ro-3306 for 75 min. Cells were treated with 5 ATRi from 15-75 min. (I) Immunoblot of MCM4 in chromatin fraction and CDC25A in the soluble fraction prepared from CD8^+^ T cells treated with ATRi and/or 5 mM HU in the sequence indicated. (J) Cartoon of a model in which ATR limits origin firing across active replicons. Green circles indicate origins that fired and red circles indicate origins that have not fired. (E-F) Mean and SD bars shown.**:P< 0.01,***:P<0.001****:P <0.0001 by two-tailed, unpaired t-test.

### ATR kinase activity limits origin firing across active replicons in CD8^+^ T cells

ATR limits origin firing across active replicons in unperturbed human cancer cells and fibroblasts (Moiseeva *et al*., 2019b). To investigate whether this mechanism is conserved in CD8^+^ T cells, we first examined the hyper-phosphorylation of chromatin bound MCM4 which is associated with activation of the replicative helicase (Sheu et al., 2016). ATRi induced a mobility shift in MCM4 in chromatin extracts prepared from proliferating CD8^+^ T cells (Figure 3C).

We then used Repli-seq to determine whether ATRi induces changes in the replication timing program. We treated proliferating CD8^+^ T cells with ATRi for 30 min and then added 5-bromo-2-deoxyuridine (BrdU) for 30 min. CD8^+^ T cells were sorted into 2N-3N and 3N-4N populations and BrdU-labelled nascent DNA was purified and sequenced. ATRi did not induce late origin firing in early S phase in proliferating CD8^+^ T cells (Figure 3D and Figure S4A and B). Furthermore, the distribution of DNA synthesis between early and late-S phase was not significantly different between proliferating CD8^+^ T cells and fibroblasts (Figure S4C).

To determine whether ATRi increased DNA synthesis, we treated proliferating CD8^+^ T cells with ATRi for 30 min and then added EdU for 30 min. ATRi increased the relative fluorophore-EdU intensity in proliferating CD8^+^ T cells in both in early S (2N-3N) and late S (3N-4N) phase cells (Figure 3E).

Next, we combed DNA purified from proliferating CD8^+^ T cells, primary fibroblasts, and B16 cells. Cells were treated with ATRi for 30 min followed by 5-iodo-2-deoxyuridine (IdU) for 10 min and 5-chloro-2-deoxyuridine (CldU) for 20 min. ATRi decreased inter-origin distance and replication track length in proliferating CD8^+^ T cells, fibroblasts, and B16 cells (Figure 3F and Figure S4D). Thus, ATRi induces dormant origin firing across active replicons in proliferating CD8^+^ T cells.

### ATR kinase activity limits CDK1-dependent RIF1 serine-2153 phosphorylation in proliferating CD8^+^ T cells

ATRi-induced origin firing in human cells is associated with a CDK1-dependent phosphorylation of RIF1 serine-2205 that disrupts an association between RIF1 and PP1 (Moiseeva *et al*., 2019b). RIF1 is a regulatory subunit of PP1 and PP1 opposes CDC7 kinase activity at replication origins (Hiraga et al., 2017; Sukackaite et al., 2017). To determine whether this mechanism is conserved in CD8^+^ T cells we treated proliferating CD8^+^ T cells with CDK1i Ro-3306, CDK2i CVT-313, and CDC7i PHA-767491 for 15 min and then ATRi for 1 h. ATRi induced a mobility shift in MCM4 in chromatin extracts prepared from proliferating CD8+ T cells and this was blocked by 5 µM Ro-3306 and 20 µM PHA-76749, but not by CVT-313 (Figure 3G).

We generated a rabbit monoclonal antibody that identifies mouse RIF1 phosphoserine-2153 (phosphoserine-2205 in human) (Figure S5A-C). ATRi induced RIF1 phosphoserine-2153 in proliferating CD8^+^ T cells and this was blocked by Ro-3306 (Figure 3H). To our knowledge, this is the first antibody that identifies mouse RIF1. Two RIF1 isoforms identified in mouse embryonic stem cells genetically engineered to express RIF1-FLAG-HA were similar in size to the proteins identified here (Sukackaite *et al*., 2017).

ATR kinase-dependent CHK1 activity phosphorylates CDC25A causing its degradation in the proteasome after DNA damage (Liu *et al*., 2000; Mailand *et al*., 2000). Since ATRi induces CDK1-dependent origin firing we reasoned that physiological (low) ATR-CHK1-CDC25A signaling may limit origin firing by limiting CDK1 activation. If this is correct, DNA damage-induced (high) ATR-CHK1 signaling should degrade CDC25A and this should reverse ATRi induced origin firing.

Accordingly, the sequence of treatment with a DNA damaging agent and ATRi should determine whether ATRi induces hyper-phosphorylation of chromatin bound MCM4. We treated proliferating CD8^+^ T cells with either ATRi or the RNR inhibitor hydroxyurea (HU) for 30 min and then added the second compound for an additional 30 min. When proliferating CD8^+^ T cells were treated with HU before ATRi, ATRi did not induce a mobility shift in MCM4 in chromatin extracts (Figure 3I). Taken together, these data suggest that ATR-CHK1-CDC25A signaling limits CDK1- and CDC7-kinase dependent origin firing across active replicons in proliferating CD8^+^ T cells (Figure 3J).

### Nucleosides rescue ATRi-induced CD8^+^ T cell death

Since ATRi increased DNA synthesis in CD8^+^ T cells, we hypothesized that dNTPs may be limiting for ATRi-induced origin firing in proliferating CD8^+^ T cells. To determine whether nucleosides rescue ATRi-induced death in proliferating CD8^+^ T cells, we used three cocktails reported to rescue genome instability: low N was 250 nM adenosine (A), cytidine (C), and guanosine (G), and thymidine (dT) (Aird et al., 2013); high N was 15 µM A, C, and G, and 6 µM dT; EmbryoMax (marketed for mouse embryonic stem cell culture) was 15 µM A, C, G, and uridine (U), and 6 µM dT (Halliwell et al., 2020). We observed a dose-dependent rescue of ATRi-induced death in proliferating CD8^+^ T cells when nucleosides were added to the tissue culture media 2 h prior to and 6 h after ATRi was added for 24 h (Figure 4A). EmbryoMax did not rescue ATRi-induced CD8^+^ T cell death as efficiently as high N suggesting that 15 µM U had a negative impact on the survival.

**Figure 4:**
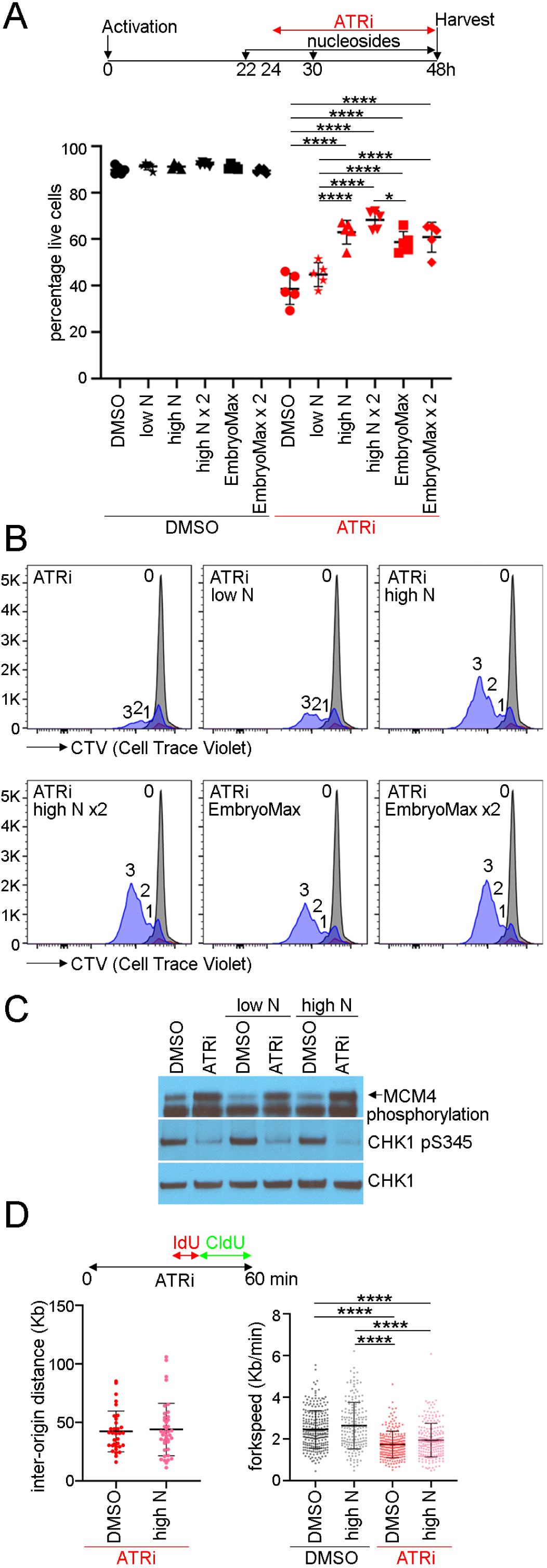
Nucleosides rescue proliferating CD8^+^ T cells from ATRi-induced cell death. (A) CD8^+^ T cells were treated with nucleoside cocktails at 22 h and 5 μM AZD6738 (ATRi) at 24 h and for x2, nucleoside cocktails at 22h and 30 h. At 48 h, live CD8^+^ T cells (eFluor 780^-^CD8^+^TCRβ^+^) was quantitated. Low N: 250 nM A, C, G, and T. High N: 15 µM A, C, G, and 6 µM dT. EmbryoMax: 15 µM A, C, G, U, and 6 µM dT. (B) CTV histograms of live (eFluor 780^-^CD44^hi^CD8^+^) cells at 24 h (grey) or 48 h (blue) treated with nucleosides and ATRi as described in panel A. Number of divisions is indicated. (C) Immunoblot of MCM4 in the chromatin fraction and CHK1 and GAPDH in the soluble fraction prepared from CD8^+^ T cells treated low N or high N for 2 h prior to ATRi for 1 h. (D) DNA combing analyses of CD8^+^ T cells treated with high N for 2 h prior to ATRi for 1 h. Cells were treated with IdU for from 30-40 min and CldU from 40-60 min during the treatment with ATRi. (A and D) (right panel) Mean and SD bars shown.*:P<0.05,****:P <0.0001 by one-way ANOVA with Tukey’s multiple comparisons. (D) (left panel): not significant by two-tailed, unpaired t-test.

dT concentrations of 1 mM or greater are used to arrest cells at the onset of S phase and this could protect proliferating CD8^+^ T cells from ATRi-induced death (Bjursell and Reichard, 1973). We did not observe cell cycle arrest in proliferating CD8^+^ T cells treated with the nucleoside cocktails used here where the maximum concentration of dT is 12 µM (Figure 4B and Figure S6A).

ATRi is a competitive ATP inhibitor and the addition of high N to the media could block the activity of the inhibitor (Foote *et al*., 2018). Addition of high N to proliferating CD8^+^ T cells 2 h prior to ATRi did not block either the ATRi-induced mobility shift in MCM4 in the chromatin fraction or the inhibition of CHK1 phosphorylation indicating that ATRi is active in the presence of high N (Figure 4C).

The addition of high N could impact ATRi-induced origin firing. Inter-origin distance and replication fork velocity in proliferating CD8^+^ T cells treated with ATRi were not changed by the addition of high N (Figure 4D). Therefore, nucleosides rescue ATRi-induced death in proliferating CD8^+^ T cells in a dose-dependent manner without impacting either the proliferation of cells, the activity of ATRi, or origin firing.

### Thymidine rescues ATRi-induced cell death

To determine whether a single nucleoside could rescue ATRi-induced death in proliferating CD8^+^ T cells, we treated with either 15 µM A, C, or G, or 6 µM dT. Addition of dT rescued ATRi-induced death in proliferating CD8^+^ T cells with similar efficacy to high N while addition of other nucleosides did not significantly affect survival (Figure 5A). We did not observe cell cycle arrest in proliferating CD8^+^ T cells treated with 6 µM dT (Figure 5B and Figure S6B).

**Figure 5:**
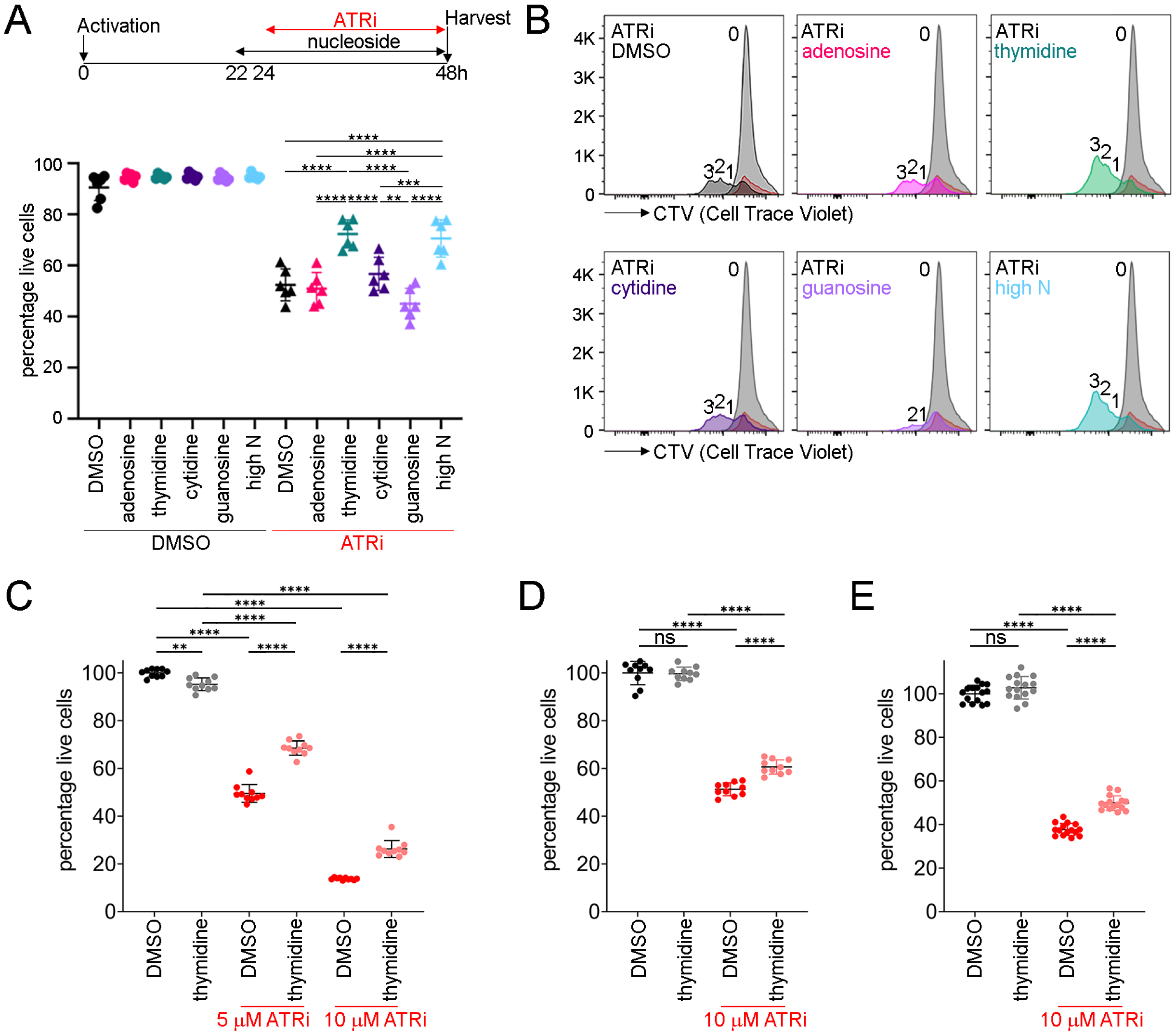
Thymidine rescues ATRi-induced cell death. (A) CD8^+^ T cells were treated with either 15 µM A, C, G, or 6 µM dT, or high N at 22 h and 5 μM AZD6738 (ATRi) at 24 h. At 48 h, live CD8^+^ T cells (eFluor 780^-^CD8^+^TCRβ^+^) was quantitated. (B) CTV histograms of live (eFluor 780^-^CD44^hi^CD8^+^) cells at 24 h (grey) or 48 h (color) treated with nucleosides and ATRi as described in panel A. Number of divisions is indicated. (C) CT26 cells were treated with 6 µM dT and 5 µM or 10 µM AZD6738 (ATRi) for 72 h and cell viability determined using CellTiter-Glo. (D-E) B16 cells were treated with 6 µM dT and 10 µM AZD6738 (ATRi) for (D) 72 h or (E) 96 h and cell viability determined using CellTiter-Glo. (A, C-E) Mean and SD bars shown. (A) **:P< 0.01,***:P<0.001,****:P <0.0001 by one-way ANOVA with Tukey’s multiple comparisons. (C-E) Data from two (C,D) or three (E) independent experiments, each with 5 biological replicates.**:P<0.01,****:P<0.0001, ns: not significant by one-way ANOVA with Sidak’s multiple comparisons (bars shown for all comparisons tested).

Next, we asked whether 6 µM dT rescued ATRi-induced death in cancer cells. We determined that 5 µM and 10 µM ATRi induced ∼50% CT26 and B16 cell death in 72 h, respectively, which was similar to that induced by 5 µM ATRi in proliferating CD8^+^ T cells (Figure 5C-E and Figure S7A). dT rescued 5 µM and 10 µM ATRi-induced death in CT26 cells at 72 h (Figure 5C). dT also rescued 10 µM ATRi-induced cell death in B16 cells at 72 h and 96 h, but the rescue was less than that observed in CD8^+^ T and CT26 cells (Figure 5D and 5E). Remarkably, dT rescues 5 µM ATRi-induced proliferating CD8+ T cell death in 24 h and CT26 cell death in 72 h with similar efficacy, and that these intervals cover ∼4 doubling times in each cell type.

### Thymidine rescues ATRi-induced, but not HU-induced cell death in proliferating CD8+ T cells

HU inhibits RNR and generates stalled replication forks that induce an intra-S phase checkpoint (Feijoo et al., 2001). To determine whether 6 µM dT rescues ATRi- and HU-induced CD8^+^ T cell death, we first determined the impact of dT, 5 µM ATRi and 5 mM HU on proliferation in CD8^+^ T cells. We treated proliferating CD8^+^ T cells with dT and 2 h later added ATRi or HU for 6 h. Cells were then replated with or without dT and allowed to recover to 48 h post-activation. Neither dT nor ATRi impacted the proliferation of CD8^+^ T cells (Figure 6A). HU for 6 h induced cell cycle arrest and this was rescued by dT during the 18 h recovery, indicating that dT is limiting for recovery from the HU-induced cell cycle checkpoints in CD8^+^ T cells.

**Figure 6:**
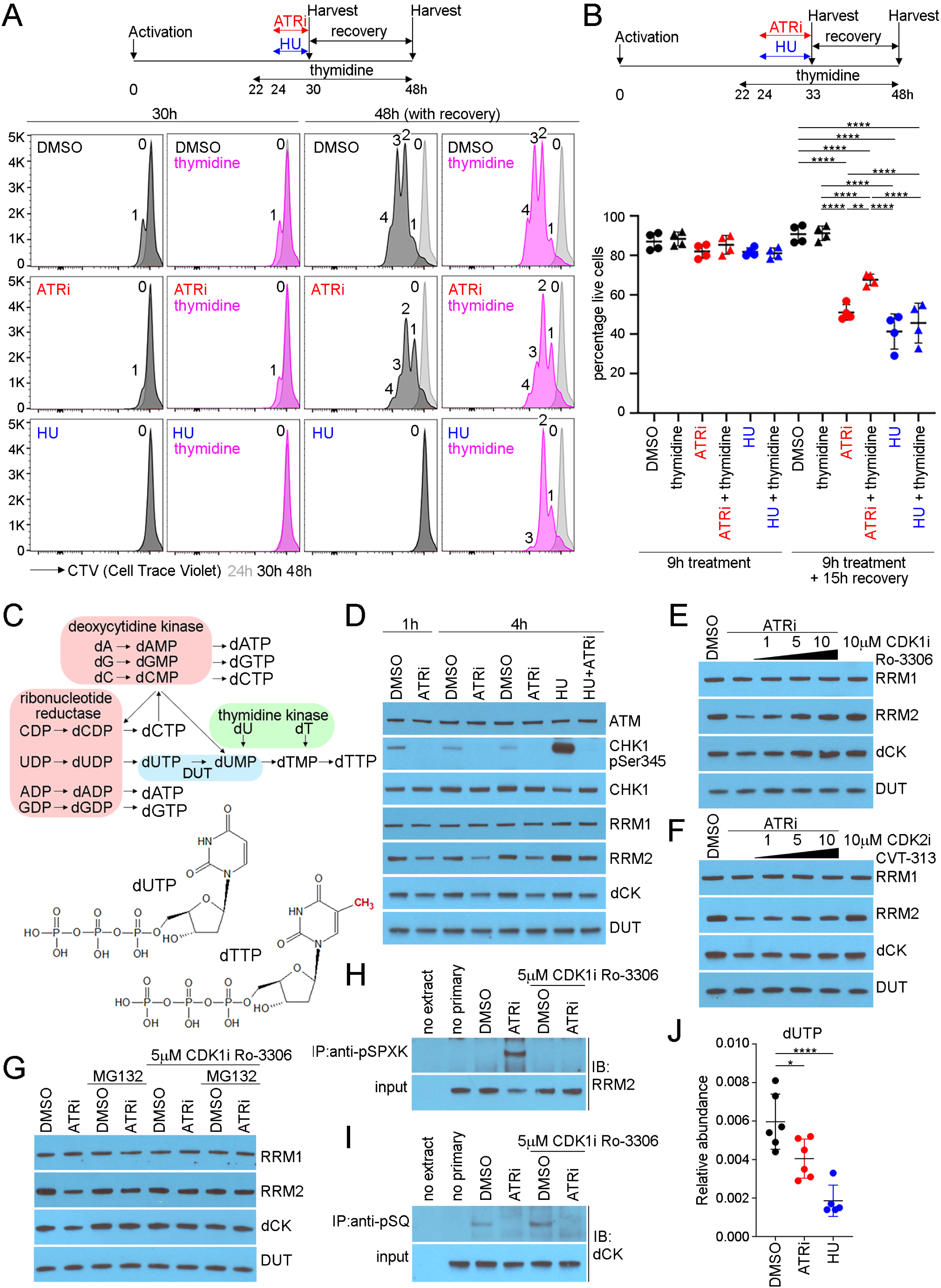
ATRi induces degradation of RRM2 and dCK in proliferating CD8^+^ T cells. (A) CD8^+^ T cells were treated with 6 µM T at 22 h and either 5 µM AZD6738 (ATRi) or 5 mM HU at 24 h. At 30 h, cells were re-plated with and without 6 µM dT in the absence of ATRi and HU. CTV histograms of live (eFluor 780^-^CD44^hi^CD8^+^) cells at 24 h (grey), and 30 h and 48 h (with recover y) for ATRi- and HU-treated (6 h) samples (DMSO treated histograms in black; thymidine treated histograms in pink). Number of divisions is indicated. (B) CD8^+^ T cells were activated and treated with 6 µM T at 22 h and either ATRi or HU at 24 h. At 33 h, cells were re-plated with and without 6 µM T. At 48 h, live CD8^+^ T cells (eFluor 780^-^ CD8^+^TCRβ^+^) was quantitated. (C) Nucleotide metabolism. (D-G) Immunoblots of proliferating CD8^+^ T whole cell extracts. (D) CD8^+^ T cells were treated with ATRi and HU for 1 or 4 h. (E) CD8^+^ T cells were treated with increasing concentrations of Ro-3306 (CDK1i) for 15 min followed by ATRi for 1 h. (F) CD8^+^ T cells were treated with increasing concentrations of CVT-313 (CDK2i) for 15 min followed by ATRi for 1 h. (G) CD8^+^ T cells were treated with 5 μM MG132 (protease inhibitor) and 5 µM Ro-3306 (CDK1i) for 15 min prior to ATRi for 2 h. (H,I) Whole cell extracts were prepared from CD8^+^ T cells treated with ATRi and CDK1i for 1 h. (H) Proteins immunopurified with anti-pSPXK antibodies were immunoblotted for RRM2. (I) Proteins immunopurified with anti-pSQ antibody were immunoblotted for dCK. (J) Quantitation of cellular dUTP in proliferating CD8^+^ T cells treated with DMSO, ATRi, or HU. (B,J) Mean and SD bars shown. (B) **:P<0.01,****:P<0.0001 by one-way ANOVA with Tukey’s multiple comparisons. (J) *:P<0.05,****:P<0.0001 by one-way ANOVA with Dunnett’s multiple comparisons.

Since 6 h incubations in 5 µM ATRi and 5 mM HU did not induce significant CD8^+^ T cell death (Figure S7B), we extended the incubation to 9 h. Cells were again replated with or without dT and allowed to recover to 48 h post-activation. A 9 h incubation in ATRi induced significant CD8^+^ T cell death and this was rescued by dT (Figure 6B and Figure S7C). In contrast, a 9 h incubation in 5 mM HU induced significant CD8^+^ T cell death, and while HU-induced cell cycle arrest was rescued by dT, HU-induced CD8^+^ T cell death was not rescued by dT. We conclude that rescue of ATRi-induced CD8^+^ T cell death by thymidine is associated with DNA synthesis.

### ATR kinase activity prevents the degradation of dCK and RRM2 in proliferating CD8+ T cells

Thymine (5’-methyluracil) is the only canonical nucleoside required for DNA replication that is not required for mRNA transcription. dUTP is an intermediate of pyrimidine nucleotide biosynthesis and the misincorporation of dU into genomic DNA in place of dT by DNA polymerases is mutagenic (Figure 6C) (Brynolf et al., 1978). The rate of dU misincorporation into genomic DNA is determined by the dUTP:dTTP ratio, which is determined in part by dCK, RNR, and DUT. Since ATR kinase inhibition and CDK1 phosphorylations has been associated with RNR and dCK degradation, we immunoblotted RRM1, RRM2, dCK, and DUT in proliferating CD8^+^ T cells treated with ATRi (Le *et al*., 2017; Smal et al., 2010).

ATRi reduced RRM2 and dCK protein levels in proliferating CD8^+^ T cells at 1 h and 4 h (Figure 6D). HU increased RRM2 protein levels in proliferating CD8^+^ T cells at 4 h and this was ATR kinase-dependent. ATRi-induced reduction in RRM2 and dCK was blocked by 5 µM and 10 µM CDK1i Ro-3306 (Figure 6E). The ATRi-induced reduction in RRM2 and dCK was not blocked by 5 µM CDK2i CVT-313 (Figure 6F). To determine whether RRM2 and dCK were degraded in the proteasome following CDK-dependent phosphorylation, we treated proliferating CD8^+^ T cells with 5 µM MG132, a proteasome inhibitor, and 5 µM Ro-3306, for 15 min prior to ATRi for 2 h. ATRi induced a CDK1-dependent reduction in RRM2 and dCK and this was MG132-dependent d (Figure 6G).

To determine whether ATRi induced a CDK1-dependent phosphorylation on RRM2, we treated proliferating CD8^+^ T cells with ATRi, with and without CDK1i, and immunoprecipitated proteins using a phospho-MAPK/CDK substrate (PXpSP or pSPXR/K) rabbit monoclonal and then immunoblotted RRM2 using rabbit polyclonal and conformation-specific HRP-conjugated antibodies. Antibodies that identify RRM2 phosphothreonine-33 (pTPPT mouse) and phosphoserine-20 (pSPLK mouse) are not available. RRM2 was immunoprecipitated from proliferating CD8^+^ T cells treated with ATRi (Figure 6H). Since the only sequence in RRM2 that matches the consensus of these antibodies is phosphoserine-20, we conclude that ATRi-induced the CDK1-dependent phosphorylation on RRM2.

To determine whether ATRi blocked the phosphorylation of dCK, we treated proliferating CD8^+^ T cells with ATRi, with and without CDK1i for 1 h, and immunoprecipitated proteins using a phospho-SQ/phospho-TQ substrate rabbit monoclonal and immunoblotted dCK using a rabbit polyclonal and a conformation-specific HRP-conjugated antibodies. Antibodies that identify dCK phosphoserine-74 (pSQ) are not available. dCK was immunoprecipitated from proliferating CD8^+^ T cells and this was blocked by ATRi (Figure 6I). Since the only sequence in dCK that matches the consensus of these antibodies is phosphoserine-74, we conclude that dCK is phosphorylated on this site by ATR in proliferating CD8^+^ T cells.

### The dUTP:dTTP ratio is ATR kinase-dependent in proliferating CD8+ T cells

We quantitated free deoxyribonucleotide concentrations in proliferating CD8^+^ T cells treated with ATRi or 5 mM HU for 1 h. The concentration of dUTP was decreased in proliferating CD8^+^ T cells treated with ATRi and HU (Figure 6J). In contrast, the concentrations of dCTP, dCDP, dCMP, dTTP, dTDP, dTMP, dUDP, and dUMP were increased in proliferating CD8^+^ T cells treated with HU, but not ATRi, consistent with HU inducing the intra-S phase checkpoint (Figure S8).

### ATR kinase activity is essential for genome stability in proliferating CD8^+^ T cells

To determine whether ATRi induced DNA damage in proliferating CD8^+^ T cells, we first quantitated γH2AX using flow cytometry. The number of γH2AX positive proliferating CD8^+^ T cells increased following ATRi and this was rescued by 6 µM dT (Figure 7A and 7B).

**Figure 7:**
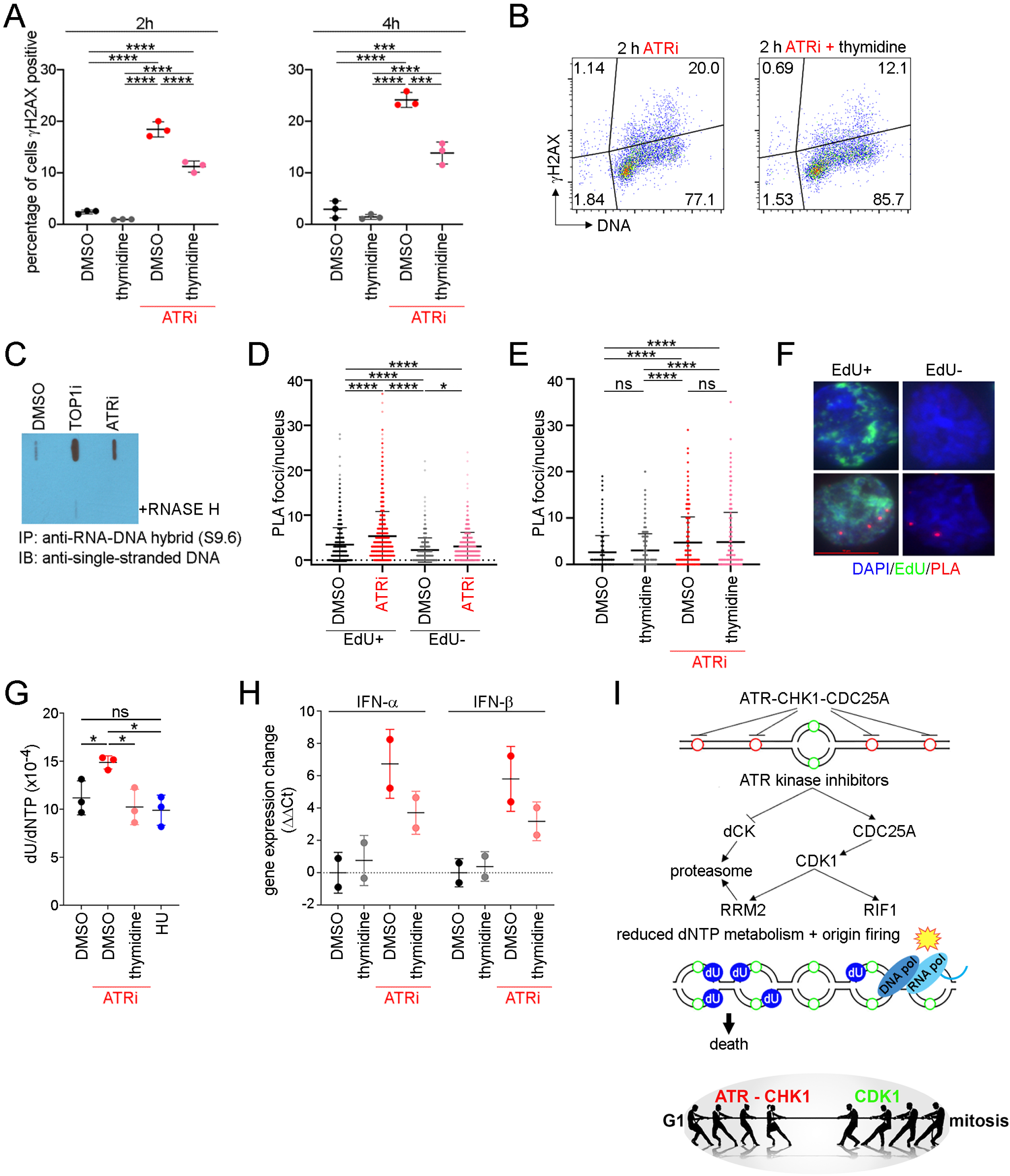
ATRi-induces genome instabilty in proliferating CD8^+^ T cells. (A) Quantitation of γH2AX-positive CD8^+^ T cells treated with 6 μM dT for 2 h followed by 5 µM AZD6738 (ATRi) for 2 h or 4 h. (B) γH2AX versus DNA content plots of CD8^+^ T cells treated with dT for 2 h followed by ATRi for 2 h. (C) CD8^+^ T cells were treated with 10 μM camptothecin (TOP1i) or ATRi for 1 h. Genomic DNA was prepared and digested with restriction endonucleases, +/-RNASE H. DNA fragments were immunopurified with anti-RNA-DNA hybrid antibody (S9.6). Purified DNA was dentured and blotted using an anti-single-stranded DNA antibody. (D-F) Proximity ligation assay (PLA) of PCNA and RNA Pol II pS5 in CD8^+^ T cells. 10 μM EdU was added during the last 15 min of the 30 min ATRi treatment. (D) Quantitation of PLA foci per nucleus in S-phase (EdU^+^) and non-S-phase (EdU^-^) CD8^+^ T cells treated with ATRi for 30 min. (E) Quantitation of PLA foci per nucleus in CD8^+^ T cells treated with dT for 2 h and then ATRi for 30 min. (F) Representative images of PLA in CD8^+^ T cells treated with ATRi for 30 min. (G) Quantitation of dU contamination in genomic DNA prepared from CD8^+^ T cells treated with ATRi for 1 h, dT for 2 h followed by ATRi for 1 h, or 5 mM HU for 1 h. (H) Quantitation of changes in IFN-α and IFN-β gene expression (ΔΔCt) in CD8^+^ T cells treated with ATRi and dT for 24 h. Each data point represents the expression change for a given biological replicate, averaged from 2 technical replicates. (I) Summary of data presented here. (A, D-E, G-H) Mean and SD bars shown. (A, D-E) *:P<0.05,***:P<0.001,****:P<0.0001, ns:not significant by one-way ANOVA with Tukey’s multiple comparisons. (G) *:P<0.05, ns:not significant by one-way ANOVA with Holm-Sidak’s multiple comparisons for DMSO vs. ATRi, DMSO vs. HU, ATRi vs. HU, and ATRi vs. ATRi plus dT.

To explore the mechanism underlying ATRi-induced γH2AX, we first hypothesized that ATRi-induced origin firing may generate R loops at sites of transcription and unprocessed Okazaki primers. We generated genomic DNA from proliferating CD8^+^ T and B16 cells treated with camptothecin (TOP1i) or ATRi for 1 h using conditions that preserve R loops and RNA primers (Chedin et al., 2021; Ginno et al., 2012). We immunopurified RNA-DNA hybrids from restriction endonuclease digested DNA using S9.6 and after extensive washing liberated the DNA from beads using proteinase K. Denatured DNA was dot blotted using an anti-single-stranded DNA antibody. RNA-DNA hybrids were increased in proliferating CD8^+^ T and B16 cells by TOP1i and ATRi (Figure 7C and Figure S9A).

Origins preferentially fire in gene promoters rather than gene coding regions (Chen et al., 2019; Petryk et al., 2016). Nevertheless, mechanisms that limit origin firing in genes to coordinate transcription and DNA replication may be critical in proliferating CD8^+^ T cells that have a short G1. We used the proximity ligation assay (PLA) to quantitate collisions between DNA replication (PCNA) and mRNA transcription (RNA polymerase II phosphoserine-5 (RNA Pol II pS5)) complexes. PCNA-RNA Pol II pS5 PLA signals increased in proliferating CD8^+^ T cells treated with ATRi for 30 min and this was not rescued by treatment with 6 µM dT (Figure 7D-F and Figure S9B). The increase in PCNA-RNA Pol II pS5PLA signals was observed in both S phase CD8^+^ T cells (EdU^+^) and, to a lesser extent, EdU^-^ CD8^+^ T cells. Proliferating CD8^+^ T cells divide extremely fast and ∼10 % of cells are anticipated to exit S phase during a 30 min treatment with ATRi.

Since ATRi-induced PCNA-Pol II collisions were not rescued by 6 µM dT, we hypothesized that the mechanism underlying ATRi-induced γH2AX might be dU contamination in genomic DNA. To quantitate DNA constituent base composition, we developed a fit-for-purpose LC-MS/MS assay that quantitated 1 N per 20,000 bases from 1 mg DNA. dU contamination in genomic DNA was increased in proliferating CD8^+^ T cells treated with ATRi for 1 h and this was rescued by treatment with 6 µM dT (Figure 7G). This observation is complementary to the significant decrease in free dUTP (Figure 6J).

Finally, since dU contamination in genomic DNA has been associated with the induction of IFN-1, we examined IFN-α and IFN-β gene expression in proliferating CD8^+^ T cells. IFN-α and IFN-β gene expression was increased in proliferating CD8^+^ T cells treated with ATRi for 24 h and this was rescued by treatment with 6 µM dT (Figure 7H).

## DISCUSSION

We investigated the impact of ATRi on mouse CD8^+^ T cells activated *ex vivo*. We show that ATRi induces dU contamination in genomic DNA, R loops, RNA-DNA polymerase collisions, IFN-1, and death in proliferating, but not naïve CD8^+^ T cells. Remarkably, we show that dT rescues ATRi-induced dU contamination, cell death, and IFN-1 expression in proliferating CD8^+^ T cells. Our data show that ATR kinase signaling integrates nucleic acid metabolism with the cell cycle to safeguard genome stability and we suggest that this signaling, rather than the ATR kinase signaling only at DNA lesions, is essential in proliferating CD8^+^ T cells.

ATR kinase activity is essential in proliferating CD8^+^ T cells activated *ex vivo*. ATRi also reduces actively proliferating CD8^+^ T cells in TIL and the periphery without impeding CD8^+^ T cell activation *in vivo*. We argue that CD8^+^ T cells activated *ex vivo* are a clinically relevant model of non-immortalized cell proliferation in which elevated dNTP and DNA synthesis are required to accommodate the replication of the mammalian genome in <4 h, and an abbreviated G1 phase is required to accommodate cell division in <6 h. After we showed that dT rescues ATRi-induced CD8^+^ T cell death in 24 h, we were able to show that dT rescues cancer cell death in 72 h. We propose that CD8^+^ T cells activated *ex vivo* are a model that may identify as yet unrealized fundamental mechanisms of DNA replication.

The ATR kinase-dependent mechanism that limits origin firing across active replicons and generates dormant origins is conserved in CD8^+^ T cells. Dormant origins are essential for genome stability and it is believed that a simple means to recover DNA replication between two stalled forks is to fire an intervening dormant origin (Shima et al., 2007). However, while the selective pressure for genome stability likely supersedes the need for rapid DNA replication and cell division in most cells, the need for elevated origin firing to accommodate rapid replication may supersede the need for genome stability in effector CD8^+^ T cells as at least 90-95% are destined to undergo cell death during contraction. We therefore reasoned that ATR kinase signaling in proliferating CD8+ T cells may limit origin firing across active replicons to integrate nucleic acid metabolism with the cell cycle to safeguard genome stability.

The ATR kinase-dependent mechanism that limits origin firing across active replicons is associated with a CDK1 kinase-dependent phosphorylation on RIF1. ATR kinase activity also limits the CDK1 kinase-dependent phosphorylation and degradation of dCK and RRM2, thereby increasing dNTP biosynthesis (Le *et al*., 2017). Furthermore, ATR phosphorylates and activates dCK (Amsailale *et al*., 2012). Accordingly, ATRi’s increase DNA synthesis while simultaneously reducing nucleotide biosynthesis. While it was known that inhibition of ATR, CHK1, and WEE1 reduces dNTP concentrations in cancer cells (Pfister et al., 2015), our findings that ATR kinase activity is essential to limit dU contamination and that dT rescues ATRi-induced CD8^+^ T cell and cancer death are, in our view, novel and unanticipated.

ATR-CHK1-CDC25A signaling is essential to restrict CDK1 activity in S phase cells and this limits origin firing and stabilizes RRM2 and *de novo* nucleotide biosynthesis (Figure 7I). We show for the first time that CDC25A protein levels are ATR kinase-dependent in unperturbed cells. This is consistent with the model proposed by Arne Lindqvist wherein DNA replication determines the timing of mitosis by restricting CDK1 activity (Lemmens et al., 2018; Lemmens and Lindqvist, 2019). It is also consistent with the finding that CDK1 activity in G2 causes RRM2 degradation (D’Angiolella *et al*., 2012). This model is attractive as DNA damage induced ATR-CHK1-CDC25A signaling in S phase would inhibit CDK1 and CDK2 and thereby origin firing and mitosis, while stabilizing RNR and *de novo* nucleotide biosynthesis. ATR would concurrently phosphorylate and activate dCK increasing salvage nucleotide biosynthesis. Thus, ATR limits and inhibits CDK1-dependent origin firing and RRM2 degradation in both undamaged and damaged cells and we suggest that CDC25A inhibition and degradation provides the switch between inhibition of origin firing across active replicons and the intra-S phase checkpoint. In the unperturbed cell cycle, CDK1 activity increases through S phase and this may ensure that all origins fire or are passively replicated before mitosis. While cyclin expression and CDC25 activities impact CDK1 activity through S, G2, and mitosis, the CDK1 and ATR phosphorylations are fundamentally the same in both in undamaged and damaged cells.

Since thymidine rescues ATRi-induced, but not HU-induced CD8+ T cell death, the underlying ATR kinase-dependent mechanism requires replicative DNA polymerase activity. HU and ATRi have different effects in proliferating CD8^+^ T cells. HU is an inhibitor of RNR that induces the ATR-CHK1-CDC25A intra-S phase checkpoint that inhibits origin firing and increases ATR kinase-dependent dCK activity. In CD8^+^ T cells treated with HU for 1 h, the concentrations of dCTP, dTTP, dCDP, dTDP, dCMP, dTMP, dUDP, and dUMP increase as DNA synthesis decreases and nucleotide salvage biosynthesis is increased. In contrast, in proliferating CD8^+^ T cells treated with ATRi for 1 h, the concentrations of these nucleotides are not changed as while DNA synthesis increases and dCK activity decreases, RNR activity is reduced, but not inhibited.

dUTP is decreased in CD8+ T cells treated with either HU or ATRi and this suggests a common underlying stress that may have increased the activity of DUT. Taken together, our data suggest that the common underlying stress is in thymidine utilization. The rate of dU misincorporation into genomic DNA is determined by the dUTP:dTTP ratio. While the mechanisms that control DUT activity are not known, it is believed that DUT hydrolyses dUTP to prevent dU contamination and provide dUMP for the *de novo* synthesis of dTTP. The concentration of dUTP may also decreased by replicative polymerases as DNA synthesis is increased in proliferating CD8^+^ T cells treated with ATRi.

dU contamination in genomic DNA is repaired by base excision repair (BER) mechanisms initiated by uracil DNA glycosylases, primarily UNG, and mismatch repair (MMR). Another potential source of dU in genomic DNA is APOBEC3A- and APOBEC3B-dependent cytosine deamination. Expression of APOBEC3A and APOBEC3B sensitives cells to ATRi, indicating a role for ATR in limiting dU in genomic DNA (Buisson et al., 2017). However, the ATRi-induced dU in genomic DNA we quantitate here is unlikely to be APOBEC3-dependent as it is rescued by dT. The removal of uracil by UNG generates an abasic site which is cleaved by APE1 to generate a single-strand break (SSBs). ATRi’s are anticipated to concentrate dU contamination at replication forks, and the concentration of dU contamination may be increased by futile cycles of BER repair synthesis if dT utilization is compromised. This could result in APE1-induced SSBs on opposite strands of the helix that generate DSBs at multiple damaged sites and cell death. This is consistent with our observation that ATRi induced γH2AX in proliferating CD8^+^ T cells.

Since the number of γH2AX positive proliferating CD8^+^ T cells treated with ATRi decreased by approximately 50% by thymidine, we reasoned that a second class of lesion might also cause cell death. Pol II can synthesize 70 bp/sec and it would, by way of example, take Pol II approximately 44 min to transcribe ATM (184,490 bp) without pausing (Darzacq et al., 2007). Since G1 is only ∼1 h in proliferating CD8^+^ T cells, transcription must be largely concomitant with DNA replication. We observed that ATRi induce R loops and transcription-DNA replication machinery collisions in proliferating CD8^+^ T cells.

ATRi induces IFN-1 expression and inflammation in the tumor microenvironment in mouse models of cancer and in the clinic when combined with radiation (Dillon *et al*., 2019; Patin et al., 2022). While the mechanism underlying these observations has not been explained, ATRi’s have been shown to increase radiation-induced IFN-1 in cancer cells and this was attributed to the generation of AT-rich DNA fragments and AU-rich RNA fragments by the combination (Feng et al., 2020). We show that ATRi induces IFN-1 in proliferating CD8^+^ T cells and that this is rescued by dT. We propose that a mechanism underlying ATRi-induced IFN-1 expression is dU contamination in genomic DNA and that this may be potentiated by radiation. We propose that ATRi-induced dU contamination contributes to dose-limiting leukocytopenia and inflammation in the clinic and CD8^+^ T cell dependent anti-tumor responses in mouse models.

Currently, there are 56 active clinical trials of ATRi’s and we propose that dU contamination in genomic DNA in proliferating immune cells contributes to the dose-limiting leukocytopenia. The chemotherapeutic and immune modulating effects of ATRi described here are important considerations for the design of schedules in the oncology clinic. Furthermore, dU contamination in genomic DNA and its subsequent repair by BER and MMR mechanisms may be confounding experiments in original research publications that describe results where cells were treated with ATRi prior to a DNA damaging agent. In many publications endpoints caused by the combination were attributed to inhibition of DNA damage signaling induced by the ATRi. Homeostatic ATR kinase signaling and the pleiotropic effects of ATRi in undamaged cells deserve consideration.

### Limitations of the study

Dynamic gradients of kinase and phosphatase activities define effective combinations of phosphorylations that drive DNA replication and the cell cycle. Our data were generated using pharmacologic kinase inhibitors and this limits our ability to separate all mechanisms underlying our observations. ATR kinase signaling is dynamic and sensitive, and ATRi induce the rapid activation of secondary signaling and feedback pathways, and this limits our ability to identify the primary pathway inhibited. Separation of the impact of ATRi on origin firing and nucleotide biosynthesis has not been possible, although we argue that we have separated the impact of ATRi on these pathways from DNA repair and mitosis. To interrogate signaling in greater depth we are generating phosphospecific antibodies that recognize RRM2 phosphothreonine-33, dCK phosphoserine-11, and dCK phosphoserine-74. In spite of these limitations, our work is impactful as we used a clinical ATRi and a primary conclusion of our studies is that the simple addition of 6 µM dT rescues ATRi-induced endpoints. As such, further studies are not limited by complex equipment and expensive reagents.

## ACKNOWLEDGEMENTS

We thank Mark O’Connor for providing AZD6738, AZD0156, AZD1775, and Olaparib.

This work was supported by R01CA236367 and R01CA204173 (CJB), R21CA259457 and DP2GM146320 (YG), R00 CA207871 (HUO), R37CA240625 (NWS and KMA), 1DP2OD024156 (GMD) and R50CA211241 from the NIH, and PRG1477 from the Estonian Research Council (TNM). This project used the Animal Facility, Cancer Pharmacokinetics and Pharmacodynamics Facility, and the Cytometry Facility that are supported in part by award P30CA047904 from the NIH.

## AUTHOR CONTRIBUTIONS

NS, FPV, AJC, PP, JJD, SSH, DP, AB, YNG, and CJB designed, performed, and analyzed experiments. HUO, NWS, and JHB designed and analyzed experiments. TNM, YW, YNG, KMA, and GMD made significant academic contributions. NS, FPV, AJC, and CJB wrote the paper.

## STAR*METHODS

### Key Resources table

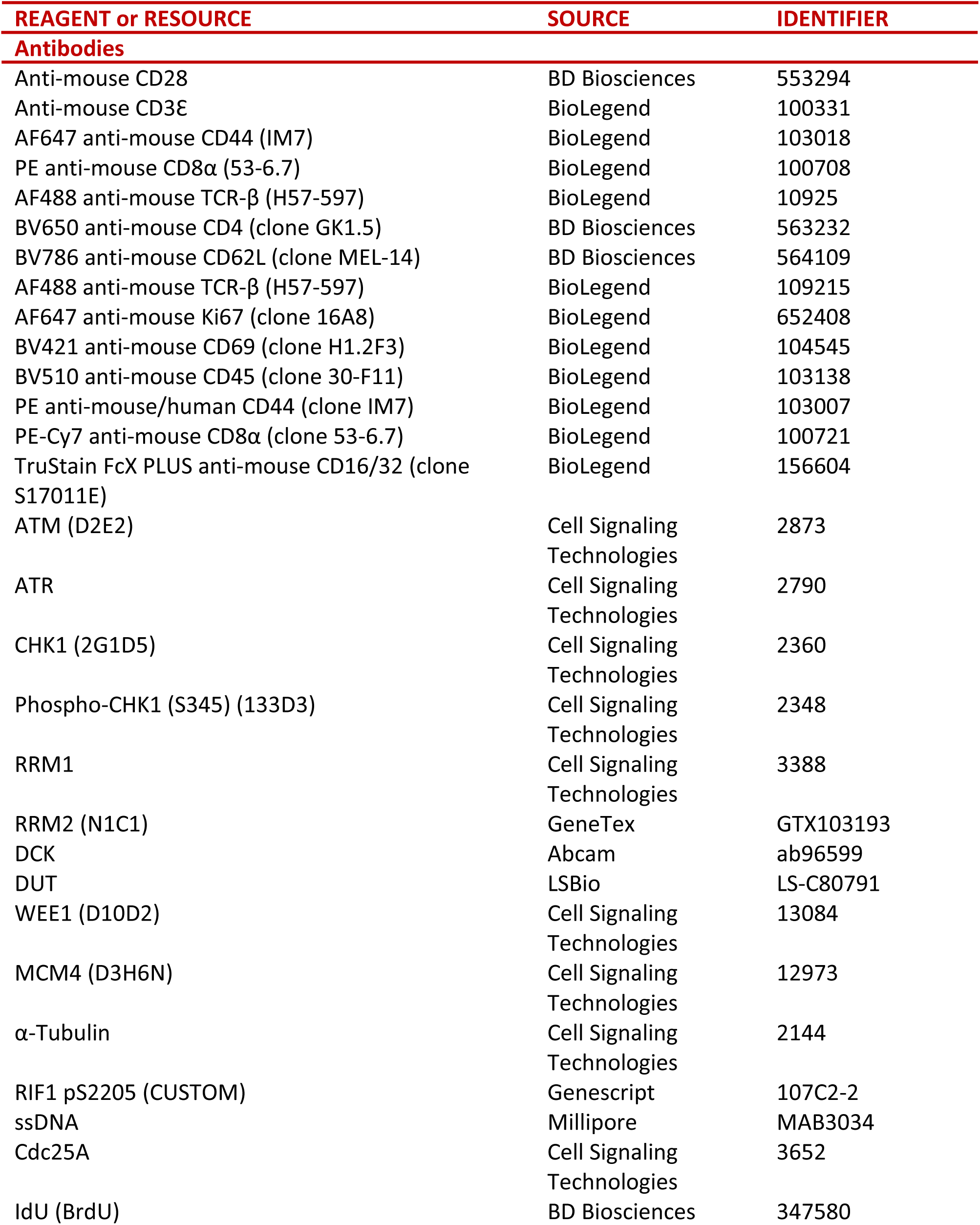
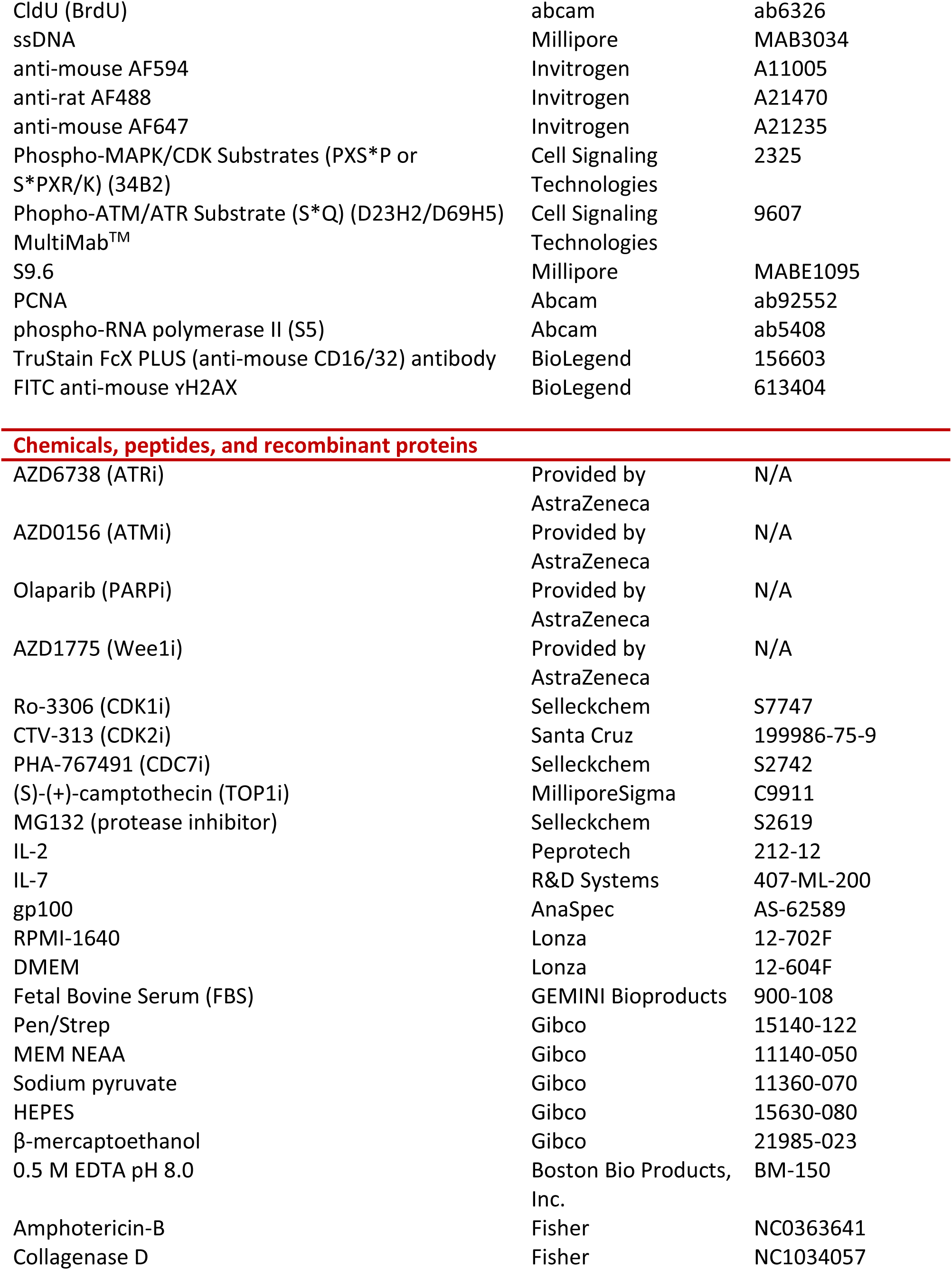
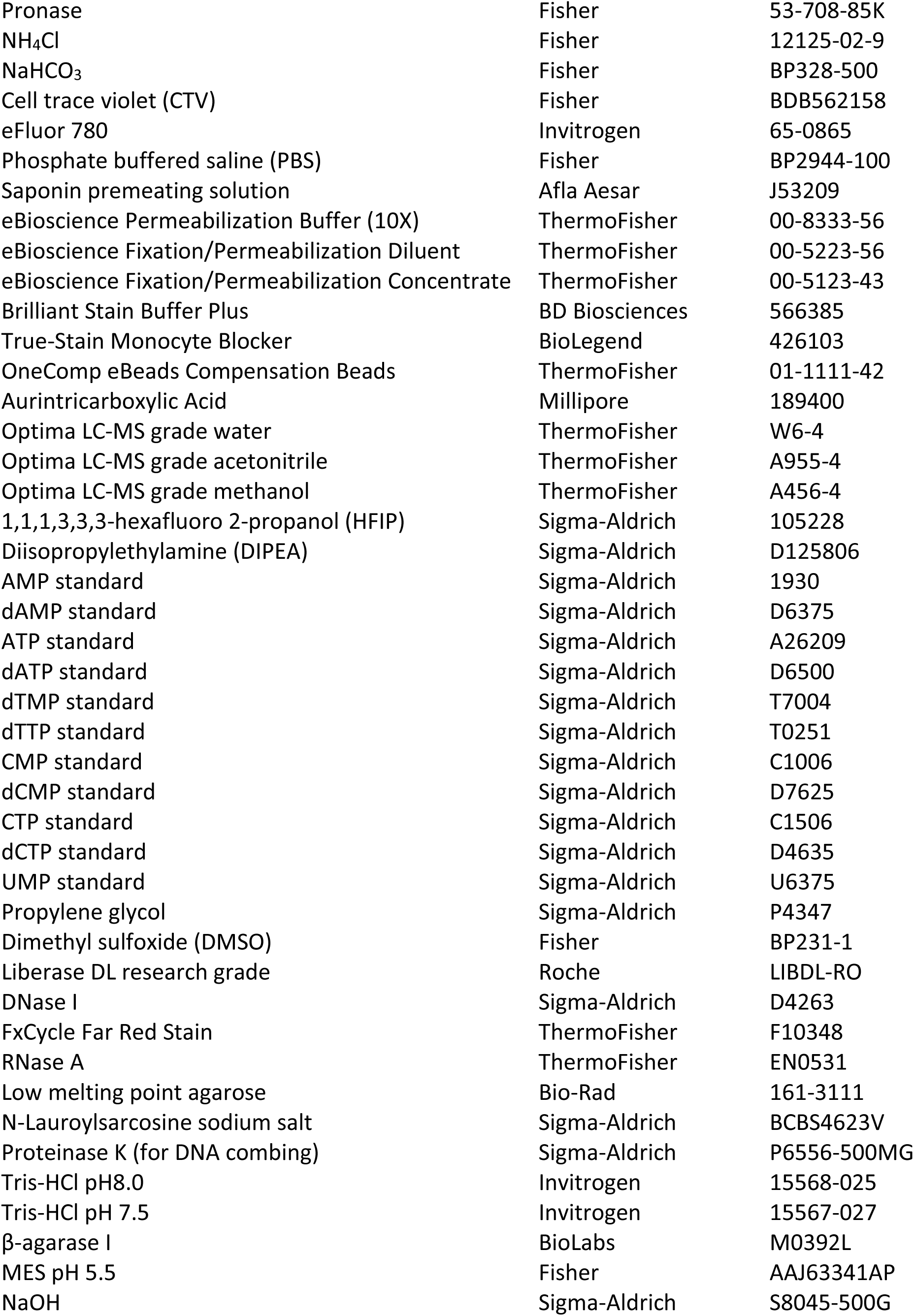
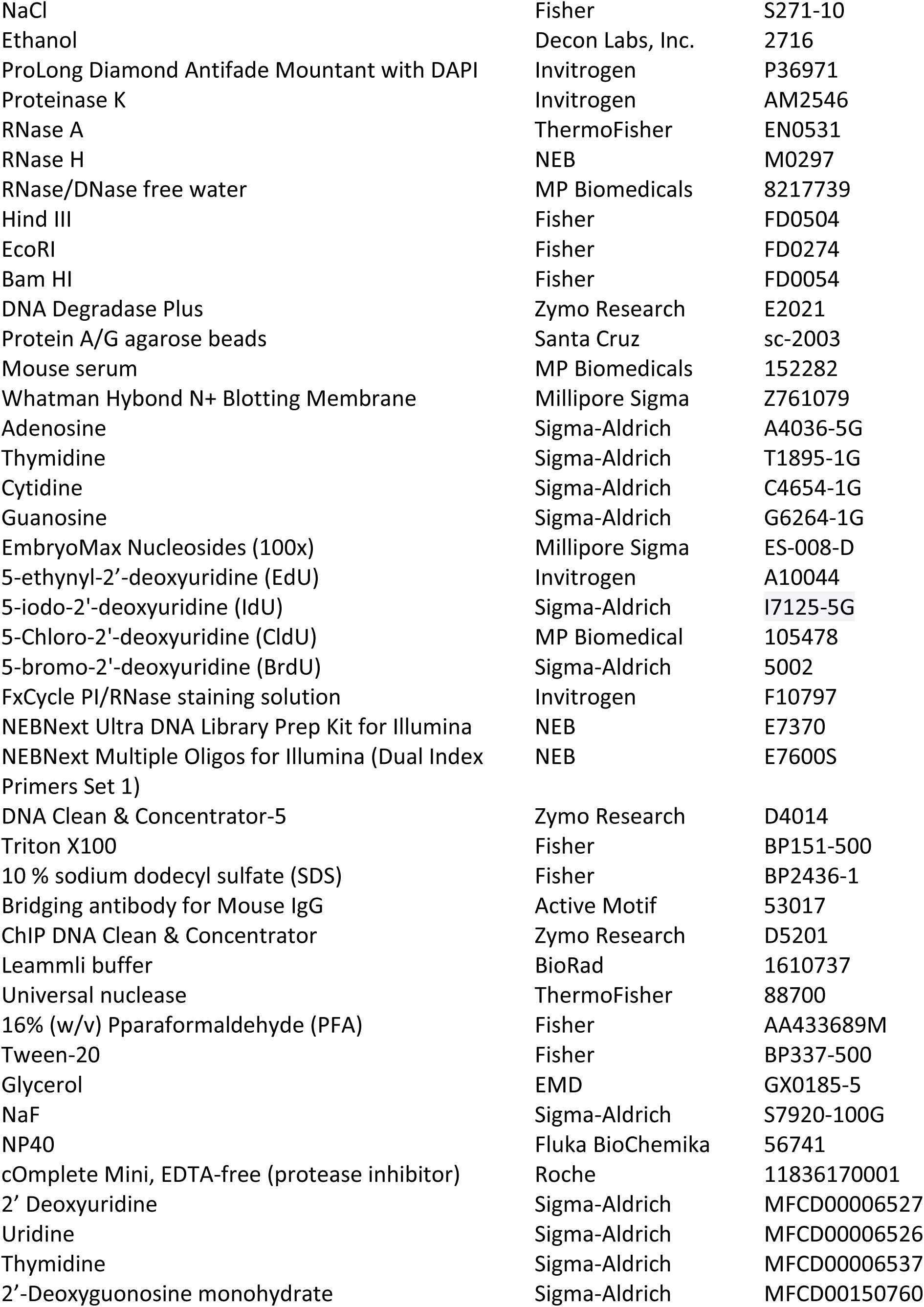
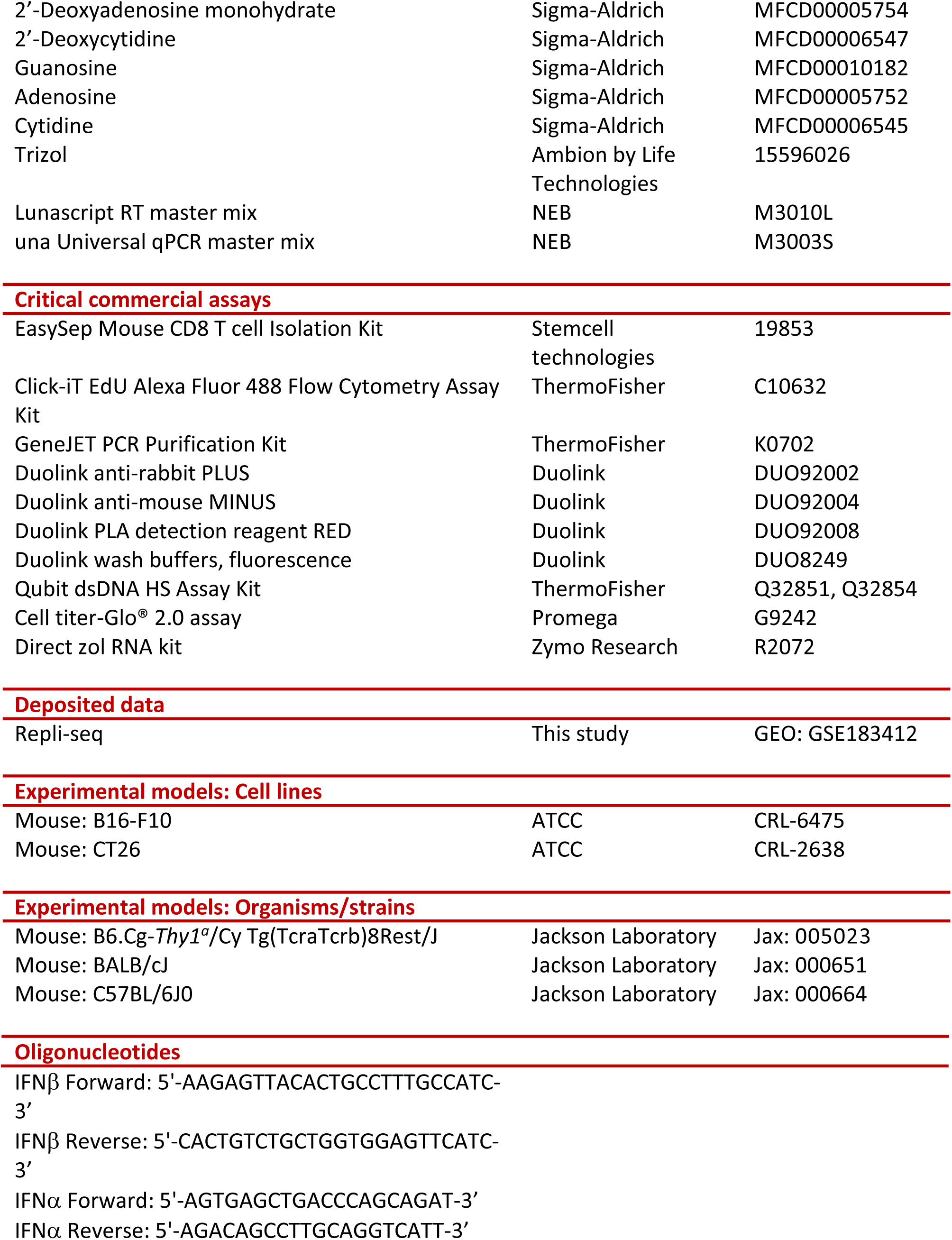
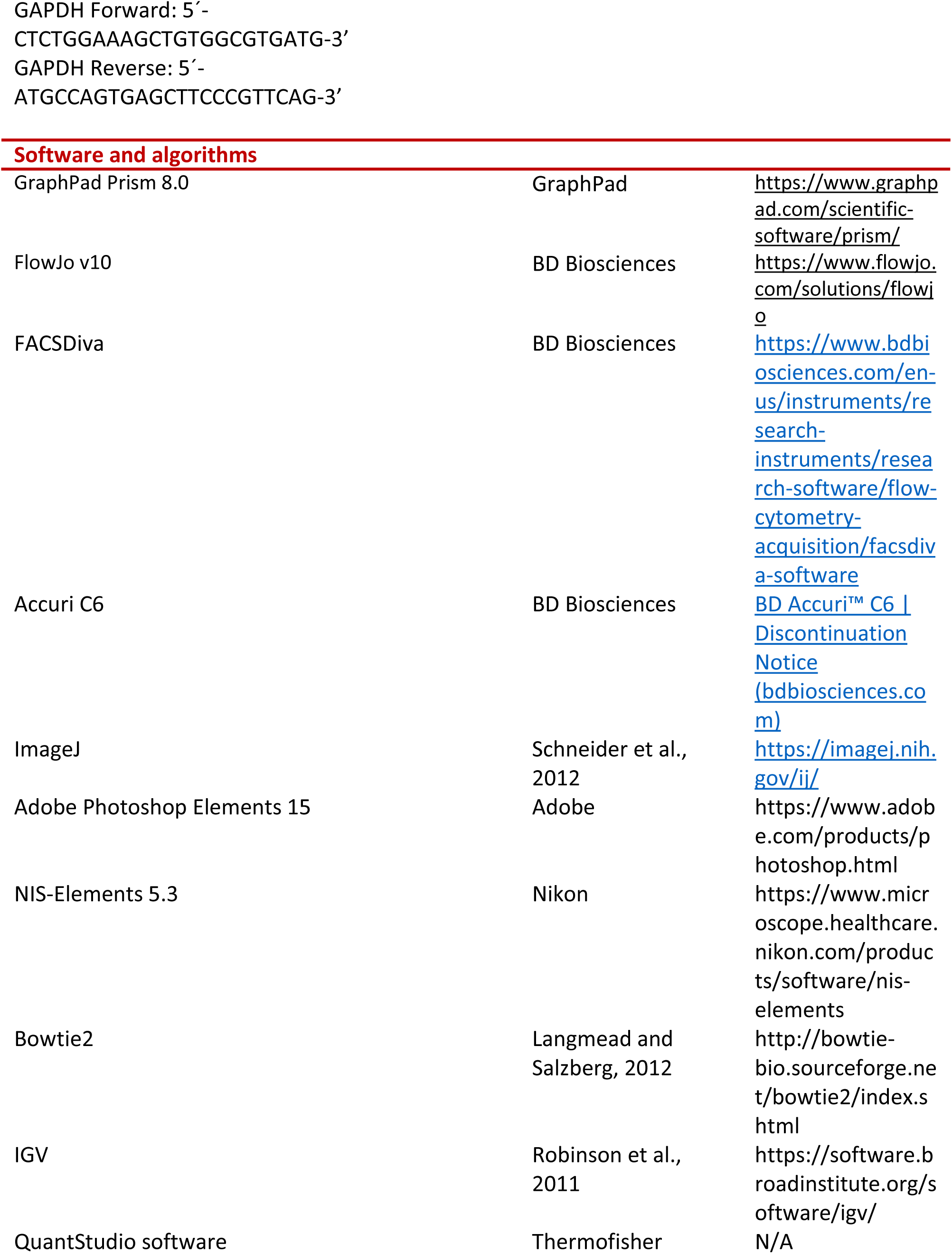
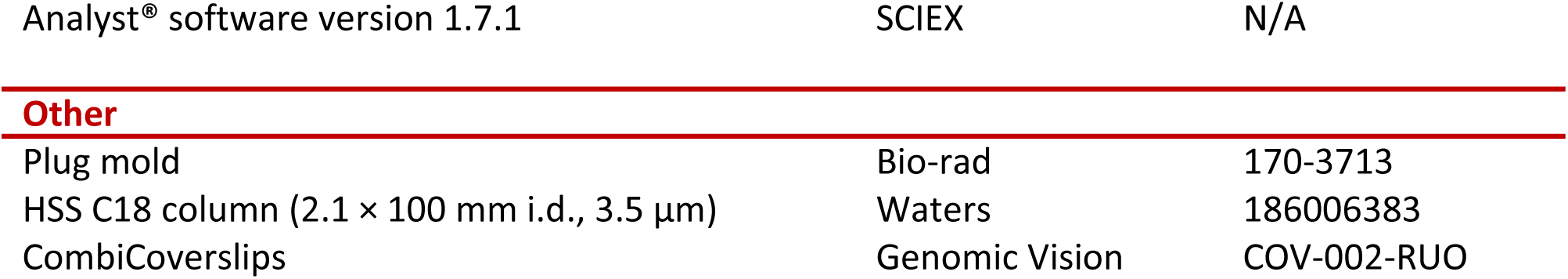

## RESOURCE AVAILABILITY

### Lead contact

Further information and requests for resources and reagents should be directed to and will be fulfilled by the lead contact, Christopher J. Bakkenist.

### Materials availability

Rabbit Monoclonal Antibody that identifies RIF1 phosphoserine-2205 (human) and RIF1 phosphorserine-2153 is available on request.

## EXPERIMENTAL MODEL AND SUBJECT DETAILS

### Animal studies

Experimental procedures were approved by the University of Pittsburgh Animal Care and Use Committees and performed in accordance with the relevant guidelines and regulations. C57BL/6, Pmel-1 TCR transgenic (B6.Cg-*Thy1^a^*/Cy Tg(TcraTcrb)8Rest/J), and female BALB/c mice were purchased from Jackson Laboratories and housed in our facility. Homozygous breeders were maintained for Pmel-1 mice. Both male and female C57BL/6 or Pmel-1 littermates were used for *ex vivo* T cell activation, age 4-8 weeks old. Female BALB/c mice were used for CT26 tumor experiments, age 8-10 weeks old.

### Cell culture

B16-F10 (ATCC CRL-6475) and CT26 (ATCC CRL-2638) were cultured in DMEM and RPMI, respectively (both Lonza), containing 10 % FBS (GEMINI Bioproducts), 100 U/mL penicillin and 100 mg/mL streptomycin (Gibco). Cells were routinely tested for mycoplasma.

## METHODS DETAILS

### Primary fibroblasts isolation and culture

Primary fibroblasts were isolated from the ears of 6-8 week Pmel-1 mice as described previously (*53*) with minor modifications. Briefly, harvested ears were first dehydrated for 5 min. in 70 % ethanol, placed in complete medium (DMEM (Lonza)containing 15 % FBS (GEMINI Bioproducts), 100 U/mL penicillin, 100 mg/mL streptomycin (Lonza), 1x MEM NEAA, 1 mM sodium pyruvate, 25 mM HEPES pH 8.0, 120 µM β-mercaptoethanol (all Gibco), and 250 ng/mL amphotericin-B (Sigma)) on a 10 cm culture dish, and then cut into 3 mm pieces. Ear pieces were placed in a 1.8 ml cryotube vial containing collagenase D-pronase solution (1.25 mg/ml pronase and 2.4 mg/ml collagenase D (both Fisher) prepared as described previously (*53*)) and incubated at 37 °C for 90 min. at 200 rpm. Digested cells were placed in a 70 μm cell strainer on a 10 cm dish with 10 ml complete medium and grinded with 10 ml syringe plunger for at least 5 min. Cells were washed 3x with complete medium and plated in a 10 cm dish containing 250 ng/ml amphotericin B (Figher). Media was replaced on the third day.

### CD8^+^ T cell isolation, activation, and culture

CD8^+^ T cells were extracted from spleen and lymph nodes of 4-8 week C57BL/6 and Pmel-1 mice. To obtain single cell suspension, spleens and lymph nodes were mechanically processed between frosted glass slides and filtered through 70 µm cell strainers (Corning). Erythrocytes were lysed in 150 mM NH_4_Cl, 10 mM NaHCO_3_, 0.1 mM EDTA pH 8.0. Pmel-1 CD8^+^ T cells were activated by incubating with R10 media (RPMI (Lonza) containing 10 % FBS (GEMINI Bioproducts), 100 U/mL penicillin and 100 mg/mL streptomycin (Lonza), 1x MEM NEAA, 1 mM sodium pyruvate, 5 mM HEPES pH 8.0, 50 µM β-mercaptoethanol (all Gibco)) supplemented with 1 µM human gp100 (25-33) (Eurogentec) and 50 U/mL IL-2 (PeproTech) for 24 h. Naïve CD8^+^ T cells were cultured in R10 media containing 5 ng/mL IL-7 (R&D systems). C57BL/6 CD8^+^ T cells were purified using the EasySep Mouse CD8 T Cell Isolation Kit (Stemcell technologies) according to the manufacturer’s instructions. C57BL/6 CD8^+^ T cells were activated by resuspending into R10 media containing 50 U/mL IL-2 (PeproTech) and 2 µg/mL anti-CD28 antibody (BD) and then plated in a culture plate pre-coated with 10 µg/mL anti-CD3_Ɛ_ antibody (Biolegend). Culture medium was exchanged to R10 containing IL-2 and the cell density was adjusted to be 0.5 – 1×10^6^ cells/ml every 24 h.

### Proliferation and survival assays

For the proliferation assay, 5×10^6^ cells/mL were stained in 8 mg/mL Cell Trace Violet (CTV, Fisher) in PBS for 10 min. Staining was quenched the addition of with 5x the volume of R10 media and then activated as above. Cells were collected on ice and surface antigens were identified using the antibodies indicated for 15 min before staining for 10 min with eFluor780 viability dye (1:4000, ThermoFisher). Cells were fixed in Fixation/Permeabilization reagent (eBioscience) and uncompensated data was collected using a LSRFortessa cytometer and FACSDiva Software (BD Biosciences). Compensation and analyses were performed using FlowJo v10 software. Gating is shown in Figure S10.

### Nuclease insoluble chromatin fractionation and immunoblotting

Purification of nuclease insoluble chromatin (NIC) and immunoblotting were performed as described previously (*23*). Briefly, cells were lysed in 50mM Tris-HCl, pH 7.5, 150mM NaCl, 50mM NaF, 0.5% Tween-20, 1% NP40, and protease inhibitors for 20 min on ice. Lysates were cleared by centrifugation, and soluble protein was mixed with 2× Laemmli buffer (BioRad) and boiled for 10 min. Pellets were washed once with nuclease-incubation buffer (150mM HEPES (pH = 7.9), 1.5mM MgCl2, 10% glycerol, 150mM potassium acetate, and protease inhibitors). Pellets were resuspended in nuclease incubation buffer containing universal nuclease for cell lysis (ThermoFisher) and were incubated for 10 min at 37 °C on the shaker. Nuclease-insoluble chromatin was pelleted by centrifugation, washed with water and dissolved in 2x Laemmli sample buffer (BioRad) and boiled for 10 min. Proteins were resolved in 4–12% Bis-Tris or 3–8% Tris-acetate gels (Life Technologies), transferred to 0.4 μm nitrocellulose membrane (Bio-Rad) and immunoblotted. Soluble fractions were blotted for the antibodies listed in Table S1 and NIC fractions were blotted for MCM4.

### *In vivo* treatments and immunophenotyping

CT26 cells (approximately 5 x 10^5^ in RPMI) were subcutaneously injected into the right hind flank of 8-10 week old mice. Treatment started 7-10 days later when tumors reached ∼60-120 mm^3^. Mice were treated daily for 3 days with 75 mg/kg AZD6738 or vehicle in a volume of 10 µL/g of bodyweight, as described previously (*7*). AZD6738 was dosed (in 50 % H_2_O, 40 % Propylene Glycol, 10 % DMSO) by oral gavage. Spleens, tumor-draining lymph nodes (DLN, right inguinal) and CT26 tumors were excised from mice at day 4. Tissues were processed to generate single cell suspensions, as described previously (*7*). Briefly, spleens and DLN were mechanically dissociated between frosted glass slides and filtered through 70 μm cell strainers (Corning). Tumors were injected in multiple locations with a total of 1.5 mL RPMI containing 50 µg/mL Liberase DL research grade (Roche) and 10 U/mL DNase I (Sigma), incubated 3 min at room temperature, cut into small pieces, incubated in a total volume of 5 mL Liberase DL/DNase solution for 15 min at 37°C, mechanically dissociated between frosted glass slides, filtered through 70 μm cell strainers (Corning), vortexed at low speed for 90 sec, and filtered again through new 70 μm cell strainers (Corning). Erythrocytes were lysed in 150 mM NH_4_Cl, 10 mM NaHCO_3_, 0.1 mM EDTA pH 8.0. for 30 sec (spleens) or 10 sec (tumors). Cells suspensions were counted using a Scepter 2.0 or 3.0 handheld counter (Millipore) and seeded at 1.5 x 10^6^ cells in 96-well round bottom plates for staining. Cells were blocked in FSC buffer (2% FBS/1x PBS) containing 0.5 µg anti-CD16/32 antibody (TruStain FcX Plus, BioLegend) for 10 min at 4°C to block non-specific binding of antibodies via Fc receptors, stained in FSC buffer containing antibodies to surface antigens, Brilliant Stain Buffer Plus (1:5, BD Biosciences), and True-Stain Monocyte Blocker (1:20, BioLegend) for 15 min at 4°C, stained with eFlour780 viability dye (1:4000, ThermoFisher) for 10 minutes at 4°C to label dead/dying cells, fixed and permeabilized in eBioscience Fixation/Permeabilization reagent (ThermoFisher) for 15 min at room temperature, and when performing nuclear staining of Ki67, stained for 45 min at room temperature in eBioscience 1x Permeabilization Buffer (ThermoFisher) containing anti-mouse Ki67 antibody. Uncompensated data were collected with a BD LSRFortessa 4-laser cytometer and BD FACSDiva software. Compensation and data analyses were performed in FlowJo V10 software. Single color compensation controls included single stained OneComp eBeads (ThermoFisher) and single stained spleen samples and matching unstained cells. Fluorescence minus one (FMO) controls were included where appropriate to empirically determine gating. Gating strategies are shown in Figure S11.

### Cell cycle analysis

For cell cycle analysis, asynchronous populations of the cells were labeled with 10 µM EdU for 30 minutes at 37 °C. Cells were harvested, washed with PBS, and fixed in cold 70 % ethanol on ice for 30 minutes. Fixed cells were permeabilized using 1x Saponin permeating solution (Alfa Aesar) diluted in 1% BSA in PBS for 15 min. EdU was identified using Click-iT EdU Alexa Fluor 488 Flow Cytometry Assay kit (Thermofisher, C10632) according to the manufacturer’s instructions. EdU-labeled cells were washed with 1x Saponin permeating solution and resuspended in 200 – 300 µl PBS containing 200 nM FxCycle^TM^ Far Red Stain (Thermofisher, F10348) and 0.1 mg/mL RNase A (Thermo Scientific, EN0531). Data were collected with a Accuri C6 cytometer and software (BD Biosciences). Data analyses were performed using Flowjo v10 software.

### Repli-seq

Library preparation was performed as described previously (*24*). Briefly, cells were treated with vehicle or 5 μM AZD6738 for total of 1 h. Cells were pulse-labeled with 10 μM BrdU (Sigma) during the last 30 min. of the treatment. Harvested cells were washed with PBS and fixed in cold 75 % ethanol. Fixed cells were washed with PBS and stained by FxCycle PI/RNase staining solution (Invitrogen) following manufacturer’s instructions. Early S and Late S fractions were sorted based on DNA content of the cells by MoFlo Astrios cell sorter (Beckman Coulter). DNA extraction was performed on cells from each fraction using PureLink Genomic DNA Mini Kit (Invitrogen) following manufacturer’s instructions. Resulting genomic DNA was sheared with the target size of 200 bp using Covaris M220 sonicator (Covaris). End-repair and A-tailing were performed using NEBNext Ultra DNA Library Prep Kit and the NEBNext Multiplex Oligos for Illumina kit (NEB) as described previously (*55*). Resulting DNA was purified using DNA Clean & Concentrator-5 (Zymo Research) following the manufacturer’s instructions. Purified DNA was denatured on a 95 °C heat block for 10 min. and cooled on ice for a few min. BrdU-labeled DNA was immunoprecipitated by incubating with the anti-BrdU antibody (BD) in the IP buffer (PBS + 0.0625 % (v/v) Triton X100) over night at 4 °C. DNA-antibody solution was mixed with protein A/G agarose beads (Santa Cruz) pre-incubated with the bridging antibody for mouse IgG (Active Motif) for 2 h at 4 °C. The mixture was washed 3x with the IP buffer followed by a TE wash. BrdU-labeled DNA was eluted by incubating the beads with the elution buffer (IP buffer + 1 % SDS) for 15 min. at 65 °C. Resulting DNA was purified by ChIP DNA Clean & Concentrator kit (Zymo Research) following the manufacturer’s instructions. Indexing and amplification were performed on purified DNA using the Multiplex Oligos for Illumina kit (NEB) as described previously (*55*). Resulting library was purified using DNA Clean & Concentrator-5 (Zymo Research) following the manufacturer’s instructions and quantified using Qubit with dsDNA HS Assay Kit (ThermoFisher) following manufacturer’s instructions. Pooled library was and sent for sequencing at the Genewiz. Sequencing was performed on an Illumina HiSeq (GENEWIZ). Raw Repli-seq reads were trimmed and filtered for quality using Trim Galore (https://www.bioinformatics.babraham.ac.uk/projects/trim_galore/). Reads were aligned using bowtie2 (*54*) against GRCm38 (mm10). Genome-wide RT profiles were constructed, scaled, and pooled for analysis, as described previously (*55*). Briefly, Log2 ratios of early versus late read counts were calculated for 50kb non-overlapping windows, Loess smoothed at 300kb windows. Data were visualized using IGV (*56*). Repli-seq data have been deposited in the Gene Expression Omnibus database (accession no. GSE183412).

### DNA combing

30 minutes post treatment, 25 µM IdU was added for 10 minutes followed by 200 µM CldU for 20 minutes (both MP Biomedical). Cells were harvested and washed twice with PBS. Cells were then resuspended in PBS, mixed with an equal volume of 1 % low melting point agarose (Bio-Rad, 161-3111), and allowed to solidify in plug molds (Bio-Rad, 170-3713). Agarose plugs were incubated in 2 mg/mL proteinase K, 50 mM EDTA, 1 % Sarkosyl, and 10 mM Tris pH 7.5 overnight at 50 °C. The buffer was replaced and the incubation was continued for an additional 6 h. Plugs were washed 5 times in TE50 (10 mM Tris-HCl pH 7.0, 50 mM EDTA) and stored at 4 °C or washed 3 more times in TE (10 mM Tris-HCl pH 8.0, 1 mM EDTA). Each plug was melted in 200 µl TE at 68 °C for 30 minutes and then digested using β-agarase I (BioLabs, M0392L) at 42 °C overnight. MES buffer, pH 5.5 was gently added to increase the volume to 2 mL and the mixture was incubated at 68 °C 30 min. DNA was combed onto CombiCoverslips^TM^ using the molecular combing system (Genomic Vision). Cover slips were baked for 2 h at 60 °C. DNA strands were denatured in 1 M NaOH and 1.5 M NaCl for 8 min, neutralized in PBS, and dehydrated with subsequent 5 minutes incubations with 70, 90, and 100 % ethanol. Halogenated nucleotides were identified with anti-IdU (1:20, BD Biosciences, 347580) and anti-CldU (1:50, Abcam, ab6326) antibodies for 1 h at 37 °C. Slides were then stained with anti-mouse Alexa 594 (1:50, Invitrogen, A11005) and anti-rat Alexa 488 (1:50, Invitrogen, A21470), followed by anti-ssDNA (1:50, Millipore, MAB3034), and then anti-mouse Alexa 647 (1:50, Invitrogen, A21235), each incubated for 30 min. Slides were dehydrated in ethanol as described above and mounted using ProLong^TM^ Diamond Antifade Mountant with DAPI (Invitrogen, P36971). DNA fiber images were acquired with a Nikon Ti inverted fluorescence microscope using NIS Elements v5.3 at 60x magnification and analyzed using Adobe Photoshop Elements 15 and ImageJ software.

### Quantification of free nucleotides by LC-HRMS

Nucleotides were analyzed from cells by LC-HRMS as previously described (*57*). Cell pellets were extracted by addition of a 50 μL (20 ng/μL) mix of all stable isotope labeled internal standards (1000 ng/sample) in 80:20 (v/v) methanol:water followed by 1 mL of −80 °C 80:20 (v/v) methanol:water before homogenization by probe tip sonication, incubation at −80 °C for 30 min, centrifugation at 17,000 rcf for 10 minutes at 4°C, evaporation of the supernatant to dryness, then resuspended in 50 μL 95:5 water: methanol. An Ultimate 3000 UHPLC equipped with an autosampler kept at 6°C and a column heater kept at 55°C using a HSS C18 column (2.1 × 100 mm i.d., 3.5 μm; Waters) was used for separations. Solvent A was 5 mM DIPEA and 200 mM HFIP and solvent B was methanol with 5 mM DIPEA 200 mM HFIP. The gradient was as follows: 100% A for 3 min at 0.18 mL/min, 100% A at 6 min with 0.2 mL/min, 98% A at 8 min with 0.2 mL/min, 86% A at 12 min with 0.2 mL/min, 40% A at 16 min and 1% A at 17.9 min-18.5 min with 0.3 mL/min then increased to 0.4 mL/min until 20 min. Flow was ramped down to 0.18 mL/min back to 100% A over a 5 min re-equilibration. For MS analysis, the UHPLC was coupled to a Q Exactive HF mass spectrometer (Thermo Scientific) equipped with a HESI II source operating in negative mode. The operating conditions were as follows: spray voltage 4000 V; vaporizer temperature 200°C; capillary temperature 350°C; S-lens 60; in-source CID 1.0 eV, resolution 60,000. The sheath gas (nitrogen) and auxiliary gas (nitrogen) pressures were 45 and 10 (arbitrary units), respectively. Single ion monitoring (SIM) windows were acquired around the [M-H]- of each analyte with a 20 m/z isolation window, 4 m/z isolation window offset, 1e6 ACG target and 80 ms IT, alternating in a Full MS scan from 70-950 m/z with 1e6 ACG, and 100 ms IT. Data was analyzed in XCalibur v4.0 and/or Tracefinder v5.1 (Thermo Scientific) using a 5 ppm window for integration of the peak area of all analytes. Standards used as calibrants and isotope labeled internal standards are indicated in the Table SM2, all were used without further purification, and no adequate commercially available diphosphate standard was found therefore the monophosphate was used as a surrogate standard.

### Quantification of nucleosides incorporated into the genome

Cells were resuspended in TE buffer containing 62.5 µg/mL proteinase K (Invitrogen, AM2546), 62.5 µg/mL RNase A (Thermo Scientific, EN0531), and 0.5 % SDS and incubated overnight at 37 °C. Genomic DNA was purified by phenol/chloroform extraction and resuspended in RNase/DNase free water. 5 µg DNA was digested with 5 µl RNase H (NEB, M0297), 3 µl Hind III (Fisher, FD0504), 3 µl EcoRI (Fisher, FD0274), and 3 µl Bam HI (Fisher, FD0054) in RNase H buffer (NEB, M0297) overnight at 37 °C. Digested DNA was purified using the GeneJET PCR Purification Kit (Thermo Scientific, K0702). DNA was further digested into single nucleosides using DNA Degradase Plus (Zymo Research, E2021). To quantitate DNA constituent base composition, a fit-for-purpose LC-MS/MS assay was implemented on a 1290 Infinity II Autosampler and Binary Pump (Agilent) and a SCIEX 6500+ triple quadrupole mass spectrometer (SCIEX). Chromatographic separation was conducted on an Inertsil ODS-3 (3 µm x 100 mm 2.1 mm) reverse phase column (GL Sciences) at ambient temperature with a gradient mobile phase of methanol and water with 0.1% formic acid. MRM transitions of all analytes and isotopic internal standards were monitored to construct calibration curves. We were able to quantitate 1 rN per 20,000 bases from 1 mg DNA.

### R-loop purification and dot blot

Genomic DNA was prepared as above. 5 µg DNA was digested with 3 µl Hind III (Fisher, FD0504), 3 µl EcoRI (Fisher, FD0274), and 3 µl Bam HI (Fisher, FD0054) +/-5 µl RNase H (NEB, M0297) in RNase H buffer (NEB, M0297) overnight at 37 °C. Digested DNA was incubated with 50 µl of protein A/G agarose beads (Santa Cruz, sc-2003) and 10 ul mouse serum (MP Biomedicals, 152282) in 1.5 mL PBS/0.5% Triton X100 for 2 h at 4 °C. Digested DNA was then incubated with 50 µl of protein A/G agarose beads and 5 µl anti-S9.6 (Millipore, MABE1095) overnight at 4 °C. Agarose beads were washed 3x for 3 min in PBS/0.5% Triton X100 binding buffer followed by PBS wash for 3 min. The beads were then incubated with 3 µl proteinase K for 2 h at 37 °C. Nucleic acids were purified by phenol/chloroform extraction and ethanol precipitation. DNA was denatured in 50 µl TE and mixed with 50 µl of 0.8 M NaOH/20 mM EDTA solution for 10 min at 95°C then placed on ice. The solution was neutralized using sodium acetate pH 7.0 and DNA sample was transferred to Whatman Hybond N+ Blotting Membrane (Millipore Sigma, Z761079). Nucleic acids were cross-linked by to the membrane by baking at 60 °C for 30 min followed by UV exposure. The membrane was blocked for 1 h with 5 % milk in TBST and blotted for anti-ssDNA (1:1000, Millipore, MAB3034).

### γH2AX analysis by flow cytometry

Cells were harvested, washed with PBS, and fixed in cold 70 % ethanol on ice for 30 minutes for immediate staining or stored at -20 °C until use. Fixed cells were washed with PBS and permeabilized using 1x Saponin permeating solution (Alfa Aesar) diluted in 1% BSA in PBS for 15 min. Cells were stained with FITC anti-mouse ʏH2AX (1:250 in FCS buffer) for 30 min. γH2AX-labeled cells were washed with 1x Saponin permeating solution and resuspended in 200 – 300 µl PBS containing 200 nM FxCycle^TM^ Far Red Stain (Thermofisher, F10348) and 0.1 mg/mL RNase A (Thermo Scientific, EN0531). Data were collected with a Accuri C6 cytometer and software (BD Biosciences). Data analyses were performed using Flowjo v10 software.

### Proximity ligation assay (PLA)

After treatment, cells were collected on glass slides using a cytospin (Thermo Scientific) at 2000 rpm for 10 min. Cells were fixed with 4 % paraformaldehyde in PBS for 15 min and permeabilized in 0.1 % Triton X-100 for 10 min. Cells were washed with PBS, blocked with the blocking solution (Duolink) for 1 h at 37 °C and incubated overnight with anti-PCNA (1:5,000, ab 92552 (AbCam)) and anti-phospho-RNA polymerase II S5 (1:100,000, ab5408 (AbCam)) antibodies in the Ab diluent solution (Duolink) at 4 °C. PLA reactions were performed using the following kits from Duolink: anti-rabbit PLUS (DUO92002), anti-mouse MUNUS (DUO92004), PLA detection reagent RED (DUO92008), and wash buffers, fluorescence (DUO8249). Slides were washed with PBS and EdU-labeled using EdU Click-It kit (Thermofisher, C10632). Cells were mounted using ProLong^TM^ Diamond Antifade Mountant with DAPI (Invitrogen, P36971). Images were acquired with a Nikon Ti inverted fluorescence microscope using NIS Elements v5.3 at 60x magnification and analyzed using Adobe Photoshop Elements 15 and ImageJ software.

### Cell-survival analysis

Cell survival analysis was performed using cell titer-Glo® 2.0 assay (Promega) according to instructions provided by the company. Briefly, tumor cells (2.5 * 10^3^/well) were seeded in 96-well microtiter plates and kept at 37°C in a 5% CO2 incubator. The next day, cells were treated with the indicated concentration of AZD6738 (an ATR inhibitor) or DMSO (control) in the presence or absence of thymidine (6 μM) for 72 or 96 h in the incubator. Next, cells were exposed to cell titer-Glo® 2.0 reagent for 10 min. at RT, and the obtained luminescence value was measured by a plate reader (BioTek Synergy/LX multimode reader). The relative cell survival analysis was calculated in relation to DMSO treated samples.

### RNA extraction and qPCR

Primary T-cells were activated with antibodies as described above and kept for expansion at 37°C and 5% CO2-incubator using IL-2 containing R-10 media. At the desired cell concentration, T-cells (10×10^6^ cells/dish) were seeded in 60 mm dish and treated with 5 μM AZD6738 in the presence or absence of thymidine (6μM) for 24h in the incubator. They were collected and centrifuged at 3000rpm for 5min. Trizol (Ambion by Life Technologies, 15596026) was added in a 1:3 ratio to the cell suspension. Total RNA was extracted using the direct zol RNA kit (Zymo Research, R2072) according to the manufacturer’s instructions. qPCR was performed on a Quantstudio3 (Thermo). Total RNA (typically 100 ng) was reverse transcribed using the Lunascript RT master mix (NEB, M3010L) and diluted 10-fold. Each PCR reaction was run as technical duplicates using 5 µl of Luna Universal qPCR master mix (NEB, M3003S), 1 µl of each 2.5 µM primer, and 3 µl of diluted cDNA. The PCR program was run according to the manufactures suggestions and quantification was performed using the quantstudio software.

Subsequent analysis of qPCR ct values was carried out in Prism (Graphpad), normalizing to GAPDH expression. The primers used are shown in the table above.

## QUANTIFICATION AND STATISTICAL ANALYSIS

### Statistical analyses

Statistical analysis was carried out using GraphPad Prism 8.0. Data in graphs show mean and standard deviation. P values <0.05 were considered significant.

## SUPPLEMENTAL FIGURES

**Figure S1.**
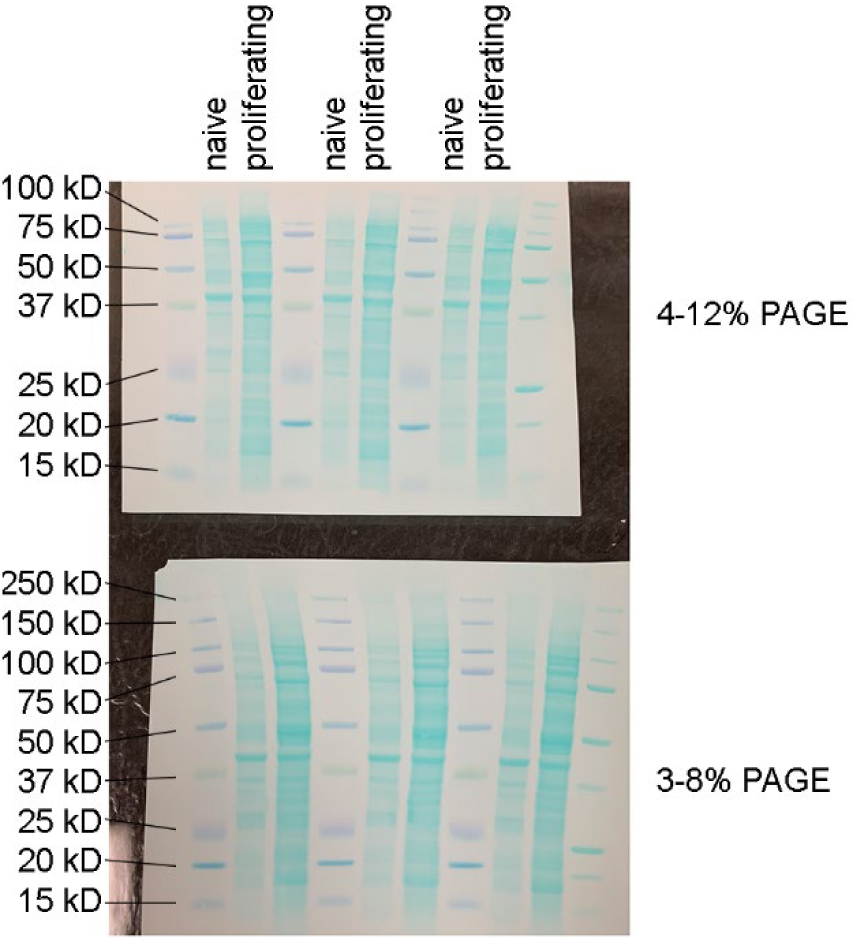
Blots of CD8^+^ T whole cell extracts. Blots of CD8+ T whole cell extracts prepared at 0 and 30 h post-activation stained with fast green prior to immunoblotting as shown in Figure 1B. The protein extracts were generated from the same number of naïve or proliferating CD8^+^ T cells.

**Figure S2.**
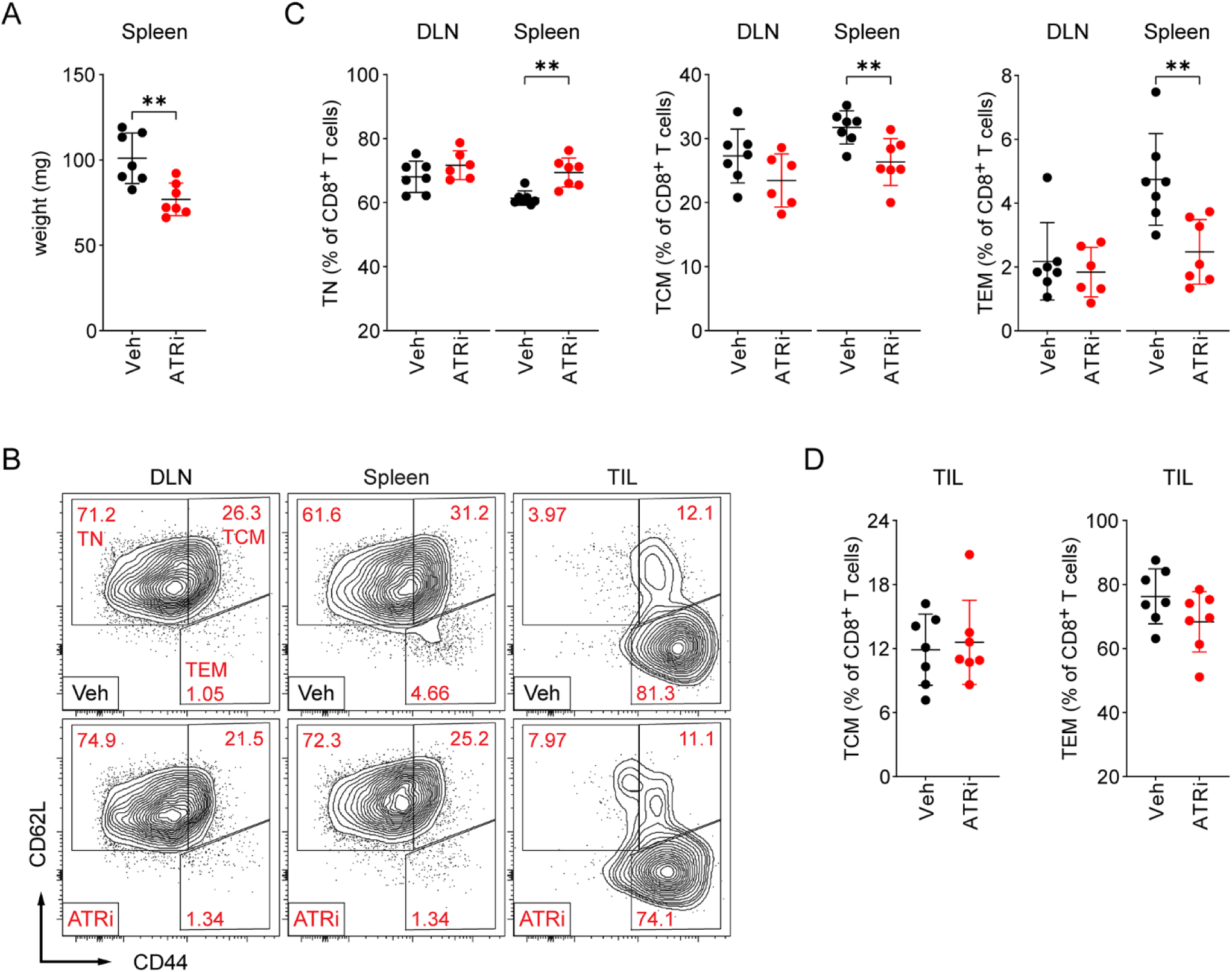
Quantitation of immune cells in mice. (A) Weight of whole spleens harvested on day 4 from CT26 tumor bearing mice treated with ATRi (75 mg/kg AZD6738) or vehicle (Veh) on days 1-3. (B-J) Immunoprofiling of tissues harvested on day 4 from CT26 tumor bearing mice treated with ATRi (75 mg/kg AZD6738) or vehicle (Veh) on days 1-3. (B) Representative contour plots of CD8^+^ T cells with naïve (TN, CD62L^hi^CD44^lo^), central memory (TCM, CD62L^hi^CD44^hi^), and effector/effector memory (TEM, CD62L^lo^CD44^hi^) phenotypes in the tumor-draining lymph node (DLN), spleen, and tumor infiltrate (TIL). (C) Quantitation of TN, TCM, and TEM CD8^+^ T cells in the DLN and spleen. **D.** Quantitation of TCM and TEM CD8^+^ T cells in the TIL. (A-D) n = 7 mice total per group (6 DLN for ATRi) from 2 independent experiments, each with 3-4 mice per group. (A, C-D) Mean and SD bars shown. **: P < 0.01 by two-tailed, unpaired t-test. Brackets not shown for comparisons that were not statistically significant.

**Figure S3.**
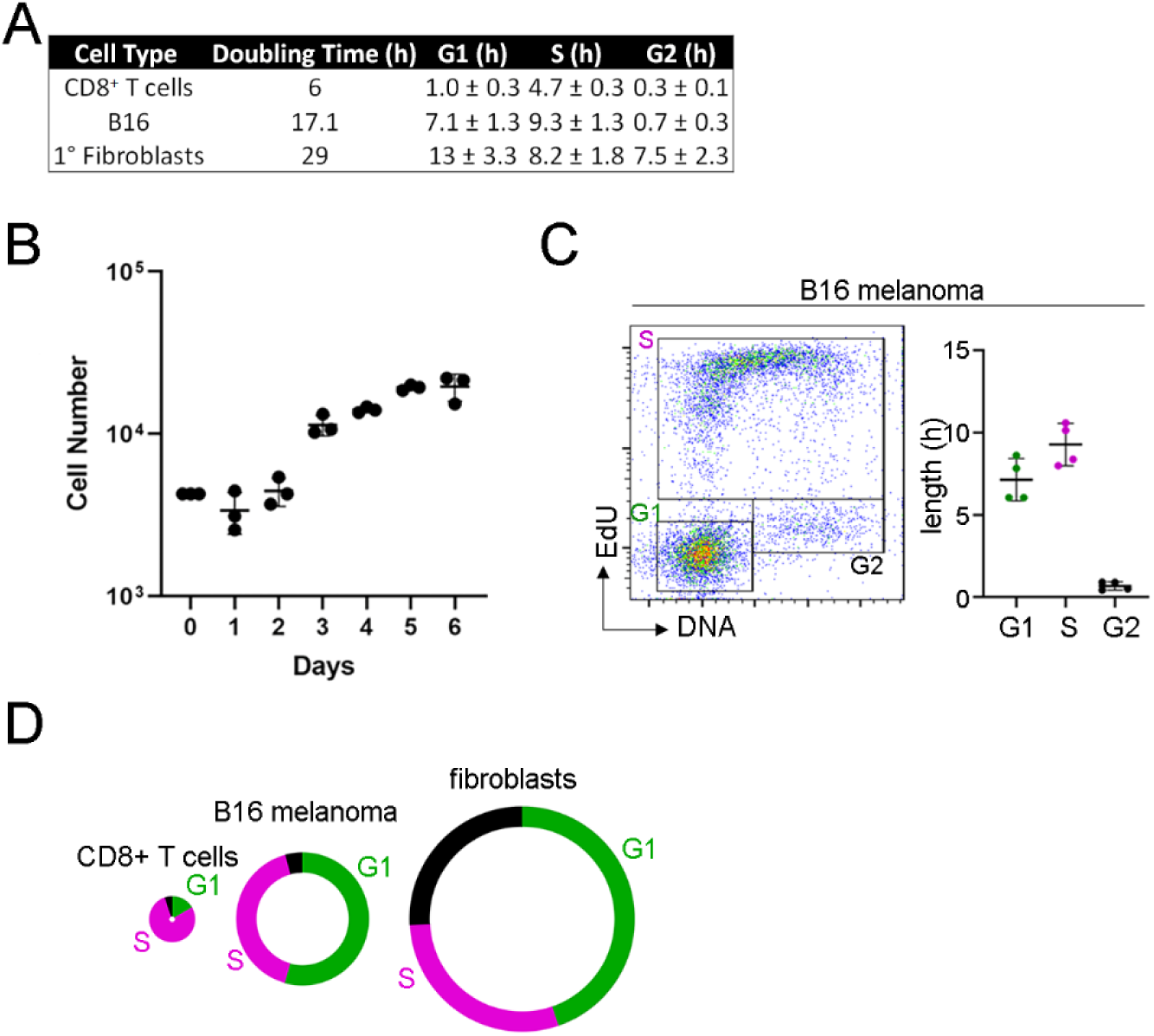
Proliferation of CD8+ T cells, B16 melanoma cells, and fibroblasts. (A) Summary of doubling times and estimated lengths of G1, S, and G2 phases in CD8^+^ T cells, B16, and primary fibroblasts. The percentage of cells in each cell cycle phase and the doubling time was used to estimate the length of G1, S, and G2/M. (B) Doubling time of B16 was previously reported (Danciu et al., 2013; Fidler, 1975) while that of primary fibroblasts was calculated with the equation (t2 – t1)/(3.32*log(n2/n1)) using linear part of the growth curve. n1 is the cell number at the time point t1 and n1 is the cell number at the time point t2. (C) EdU versus DNA histograms of B16. Mean and SD bars shown. (D) The circumference of the circle represents the doubling time and the length of G1, S, and G2/M is drawn to scale.

---

**Figure S4.**
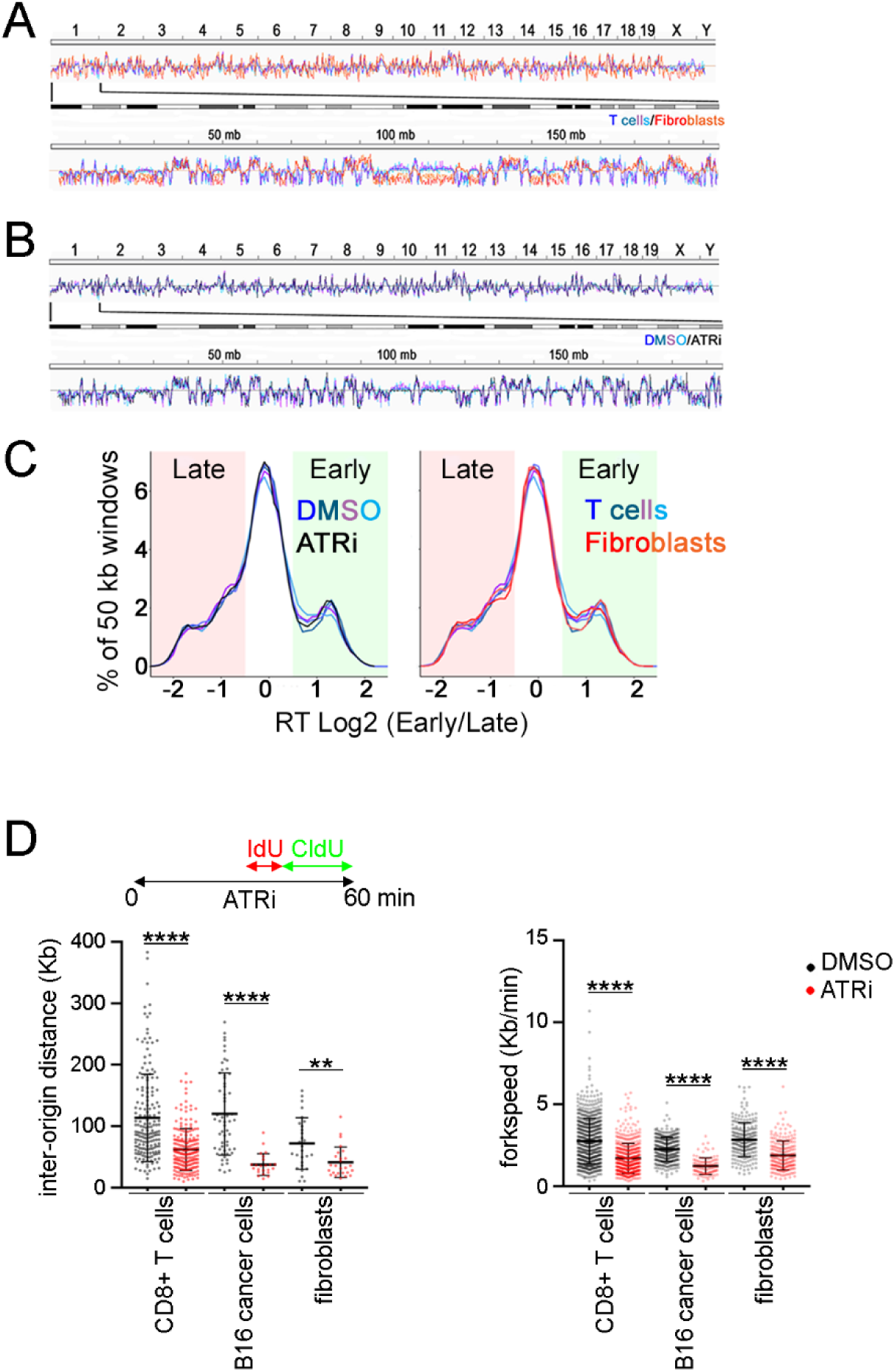
Analyses of origin firing in CD8+ T cells, B16 melanoma cells, and fibroblasts. (A) Repli-seq analyses of proliferating CD8^+^ T cells (blues) and primary fibroblasts (reds). Upper – whole genome. Lower - chromosome 1. Sequencing data are deposited at GEO (code: GSE183412). (B) Repli-seq analyses of proliferating CD8^+^ T cells treated with DMSO (blues) or 5 µM AZD6738 (black) for 1 h. (C) Genome-wide distribution of the replication timing of 50 kb genomic windows determined by repli-seq. Comparison of proliferating CD8^+^ T cells treated with DMSO (blues) or 5 µM AZD6738 (black) (left panel) or proliferating CD8^+^ T cells treated with DMSO (blues) and primary fibroblasts treated with DMSO (reds) (right panel). (D) DNA combing analyses of CD8^+^ T cells, B16 cancer cells, and fibroblasts treated with ATRi for 1 h. Cells were treated with IdU from 30-40 min and CldU from 40-60 min of the treatment.

**Figure S5.**
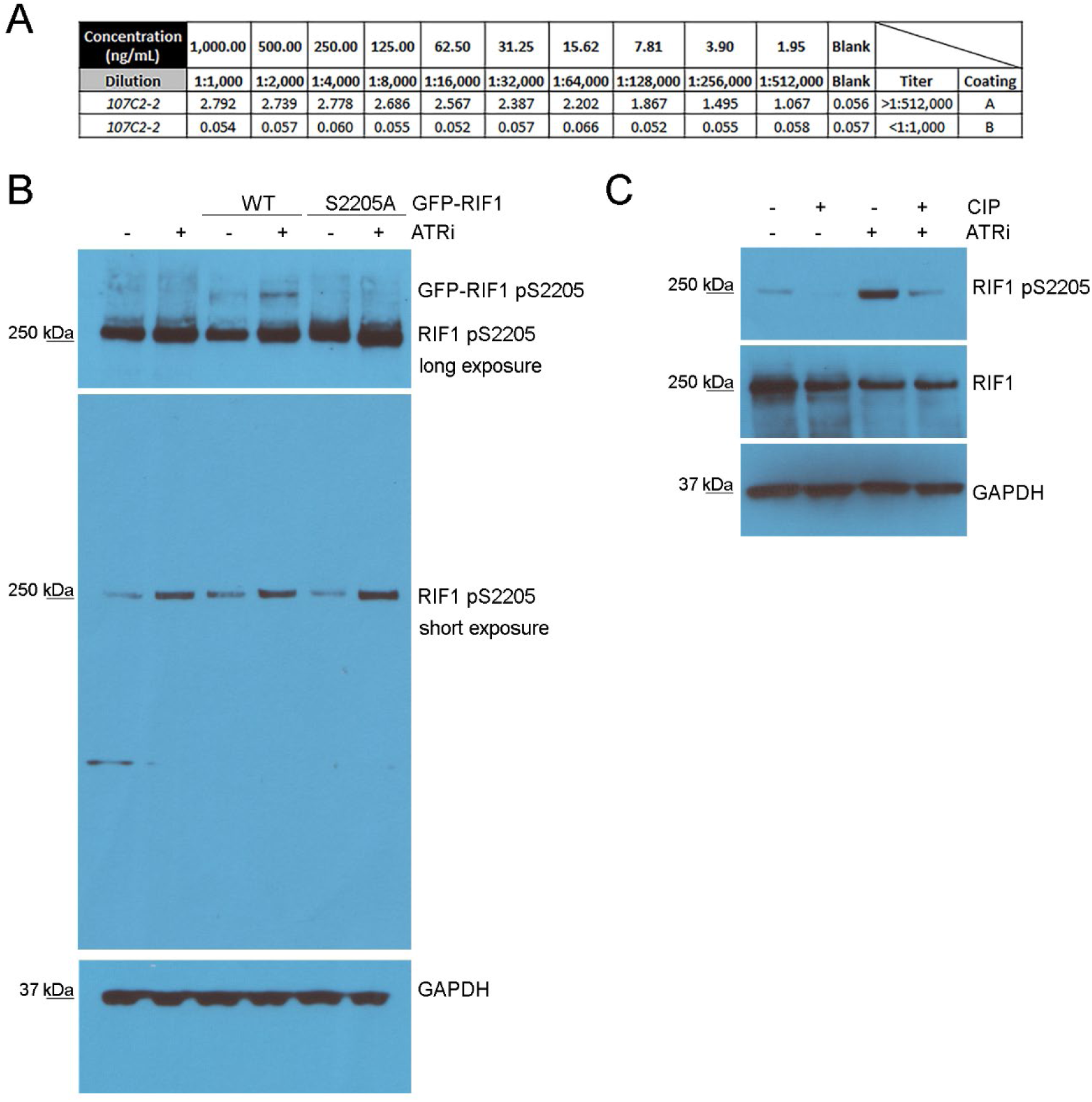
Rabbit monoclonal antibody was generated against a synthetic peptide CKVRRV(pSer)FADPI. (A) ELISA of the hybridoma supernatant shows selectivity against phosphopeptide A: CKVRRV(pSer)FADPI vs. nonphosphopeptide B: CKVRRVSFADPI. (B) GFP-RIF1 WT (wild-type) or GFP-RIF1 S2205A mutated were expressed in 293T cells. Cells were treated with ATRi for 1 h and whole cell extracts were generated and immunoblotted using the purified rabbit monoclonal antibody from A. Upper panel is a long exposure that shows that GFP-RIF1 WT, but not GFP-RIF1 S2205A mutated, is recognized by the rabbit monoclonal antibody in cells treated with ATRi. Middle panel is a typical exposure that shows that endogenous RIF1 is recognized by the rabbit monoclonal antibody in whole cell extracts of 293T cells treated with ATRi. (C) Whole cell extracts of 293T cells treated with vehicle or ATRi were treated with calf intestinal phosphatase (CIP). RIF1 is recognized by the rabbit monoclonal antibody in whole cell extracts of 293T cells treated with ATRi and this is reversed by treatment with CIP.

**Figure S6.**
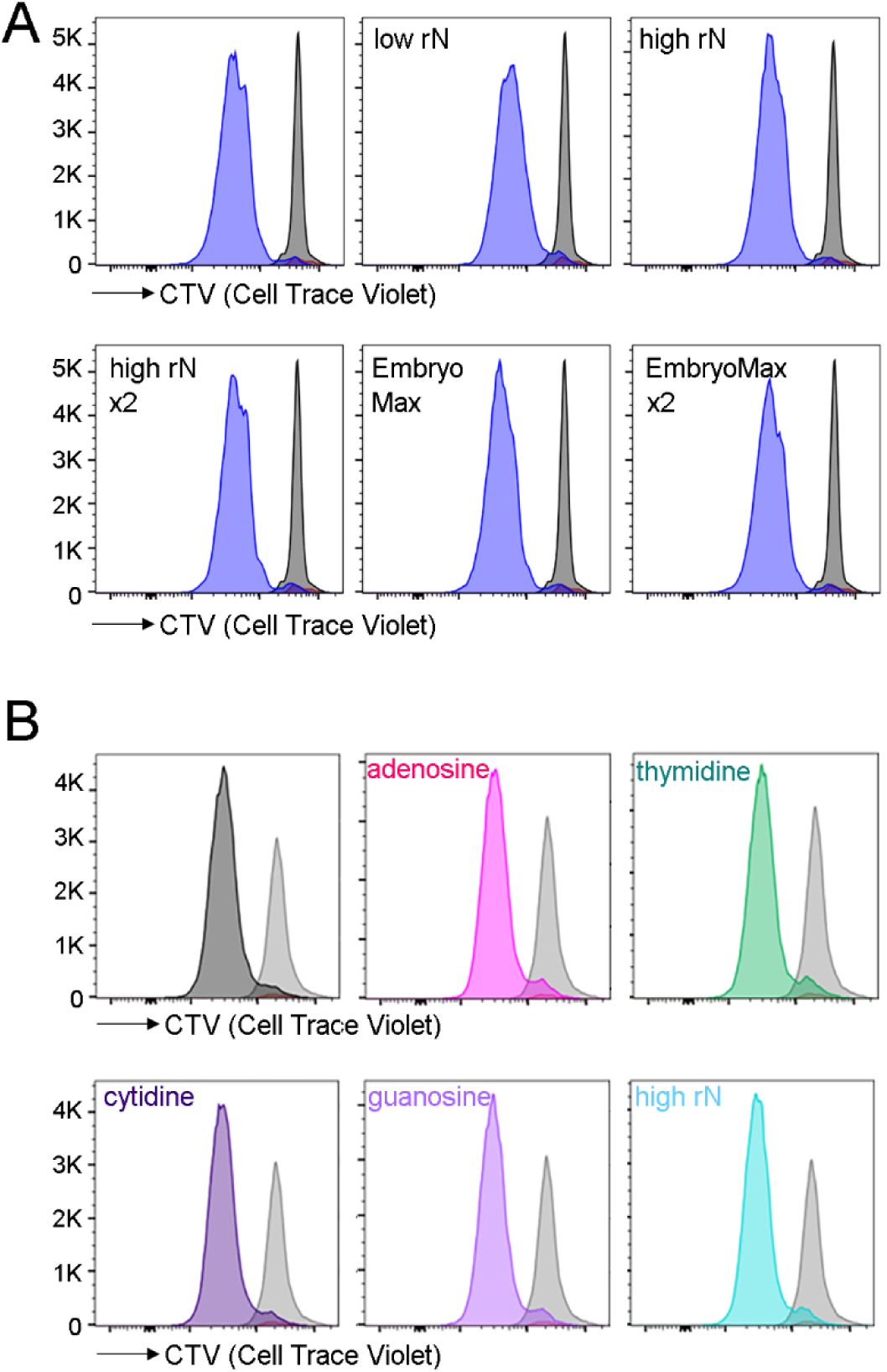
Cell proliferation in CD8+ T cells treated with nucleosides. (A) Representative overlays of CTV (proliferation) histograms for proliferating CD8^+^ T cells treated with DMSO and different nucleosides analyzed at 24 h (gray) or 48 h (blue). This is from the experiment shown in Figure 4A. Red histogram represent unactivated samples. CTV histograms are within live, CD44^hi^CD8^+^ gates. (B) Representative overlays of CTV (proliferation) histograms for proliferating CD8^+^ T cells treated with DMSO and different nucleosides analyzed at 24 h (gray) or 48 h (blue). This is from the experiment shown in Figure 5A. Red histogram represent unactivated samples. CTV histograms are within live, CD44^hi^CD8^+^ gates.

**Figure S7.**
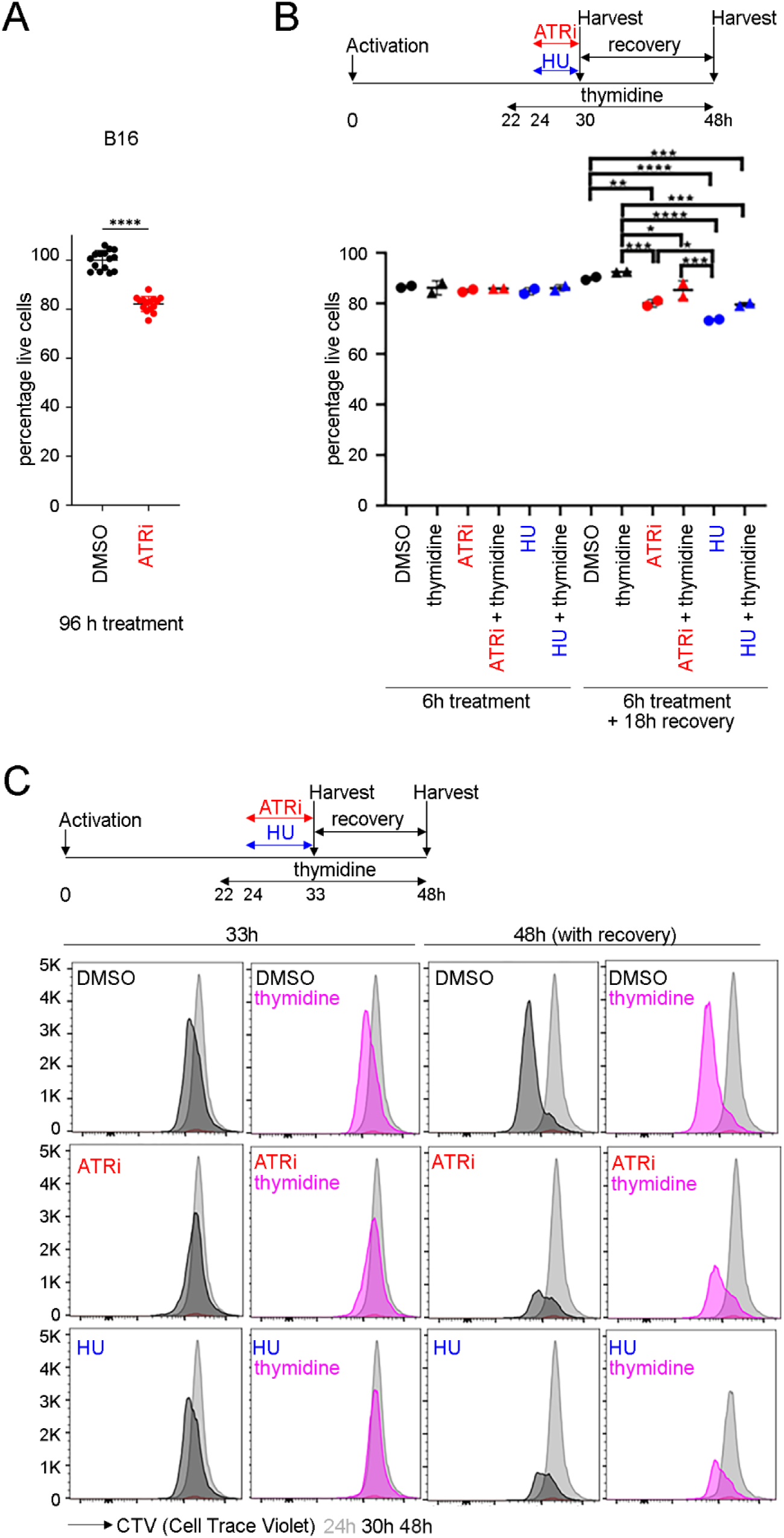
Proliferation and survival of CD8+ cells treated with ATRi, HU, and thymidine. (A) Percentage of live CD8^+^ T cells (eFluor 780^-^CD8^+^TCRβ^+^) was quantitated for the experiment shown in Figure 6A. Mean and SD bars shown. Statistics represent one-way ANOVA with Tukey’s multiple comparisons where *: P < 0.05, **: P < 0.01, ***: P <0.001, ****: P <0.0001. Brackets not shown for comparisons that were not statistically significant and between DMSO and ATRi treated samples for clarity. (B) Representative overlays of CTV (proliferation) histograms at 24 h (gray), 33 h or 48 h post stimulation for proliferating CD8^+^ T cells treated with ATRi or HU and DMSO (black) or thymidine (pink) for the experiment shown in Figure 6B. CTV histograms are within live, CD44^hi^CD8^+^ gates. .

**Figure S8.**
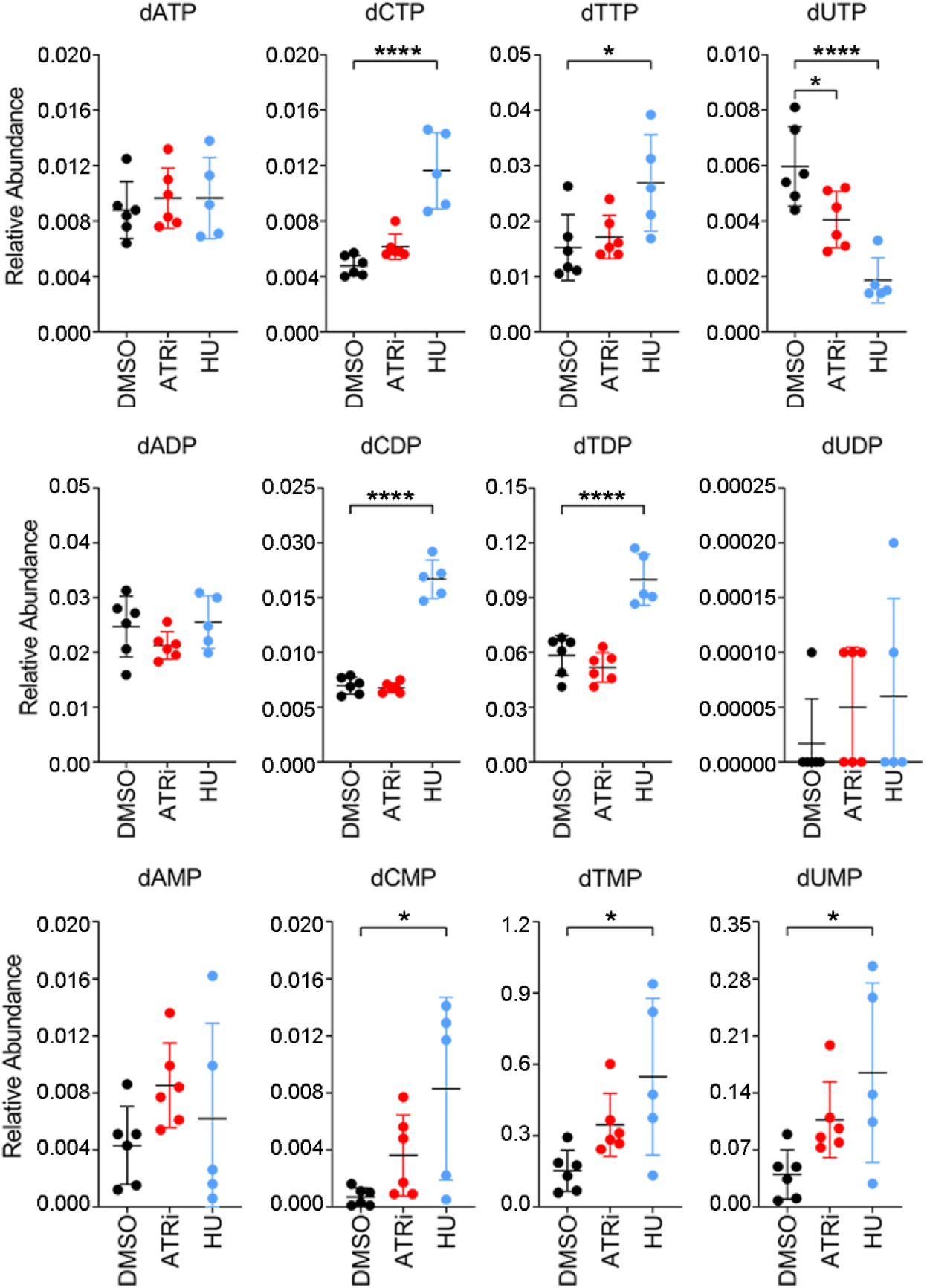
Quantitation of cellular deoxyribonucleosides in proliferating CD8+ T cells treated with DMSO, 5 µM ATRi, or 5 mM HU.

**Figure S9.**
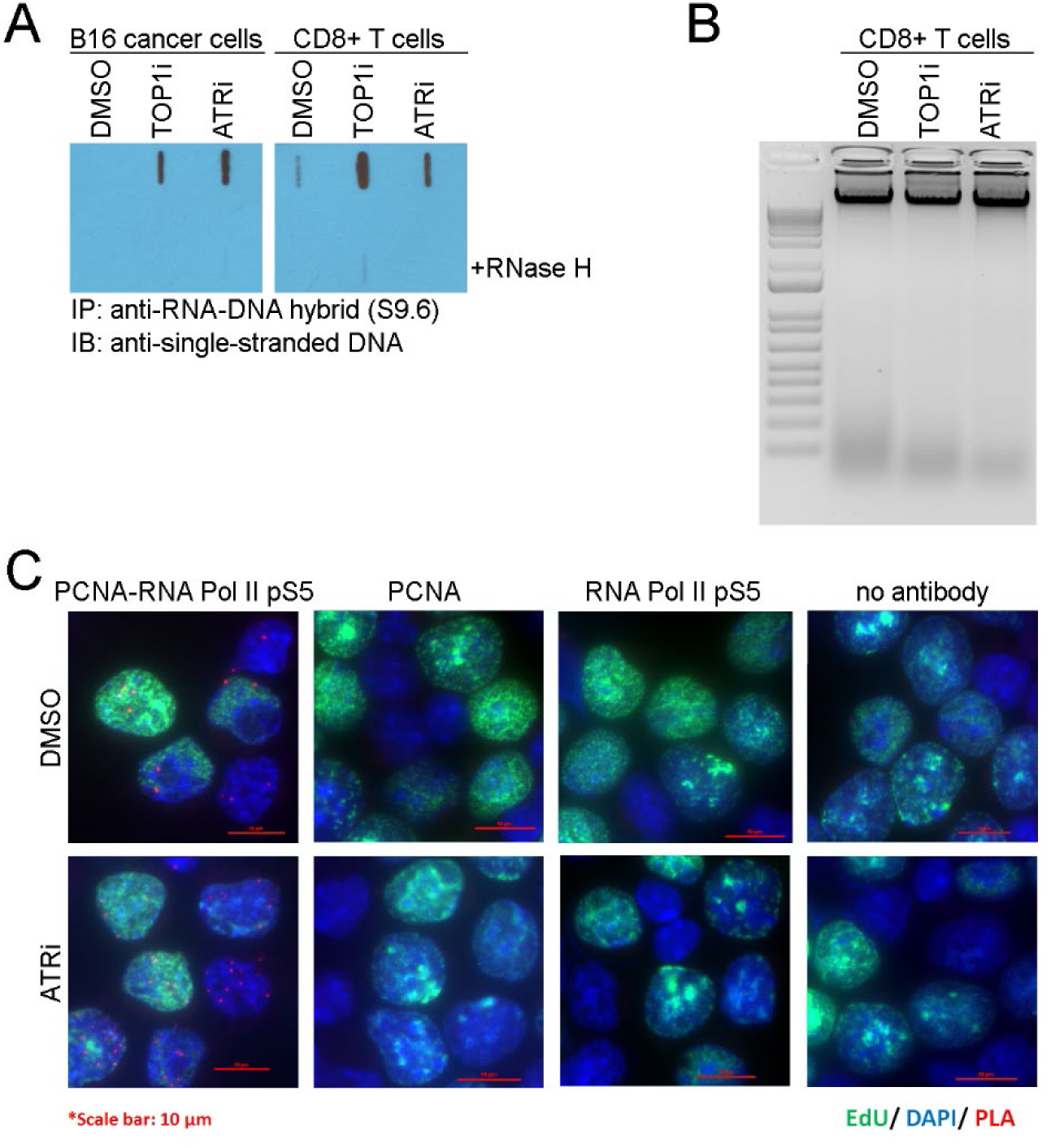
ATRi-induced R loops and PCN-RNA Pol II pS5 collisions. (A) B16 and proliferating CD8^+^ T cells were treated with 10 μM camptothecin (TOP1i) or 5 µM ATRi for 1 h. Genomic DNA was prepared and digested with restriction endonucleases, +/-RNASE H. DNA fragments were immunopurified with anti-RNA-DNA hybrid antibody (S9.6). Immunopurified DNA was denatured and dot blotted using an anti-single stranded DNA antibody. (B) Genomic DNA prepared from proliferating CD8^+^ T cells that was digested with restriction endonucleases, +/-RNASE H in Figure S8A. **C.** Proximity ligation assay (PLA) of PCNA and RNA Pol II phosphoserine-5 in proliferating CD8^+^ T cells.

**Figure S10.**
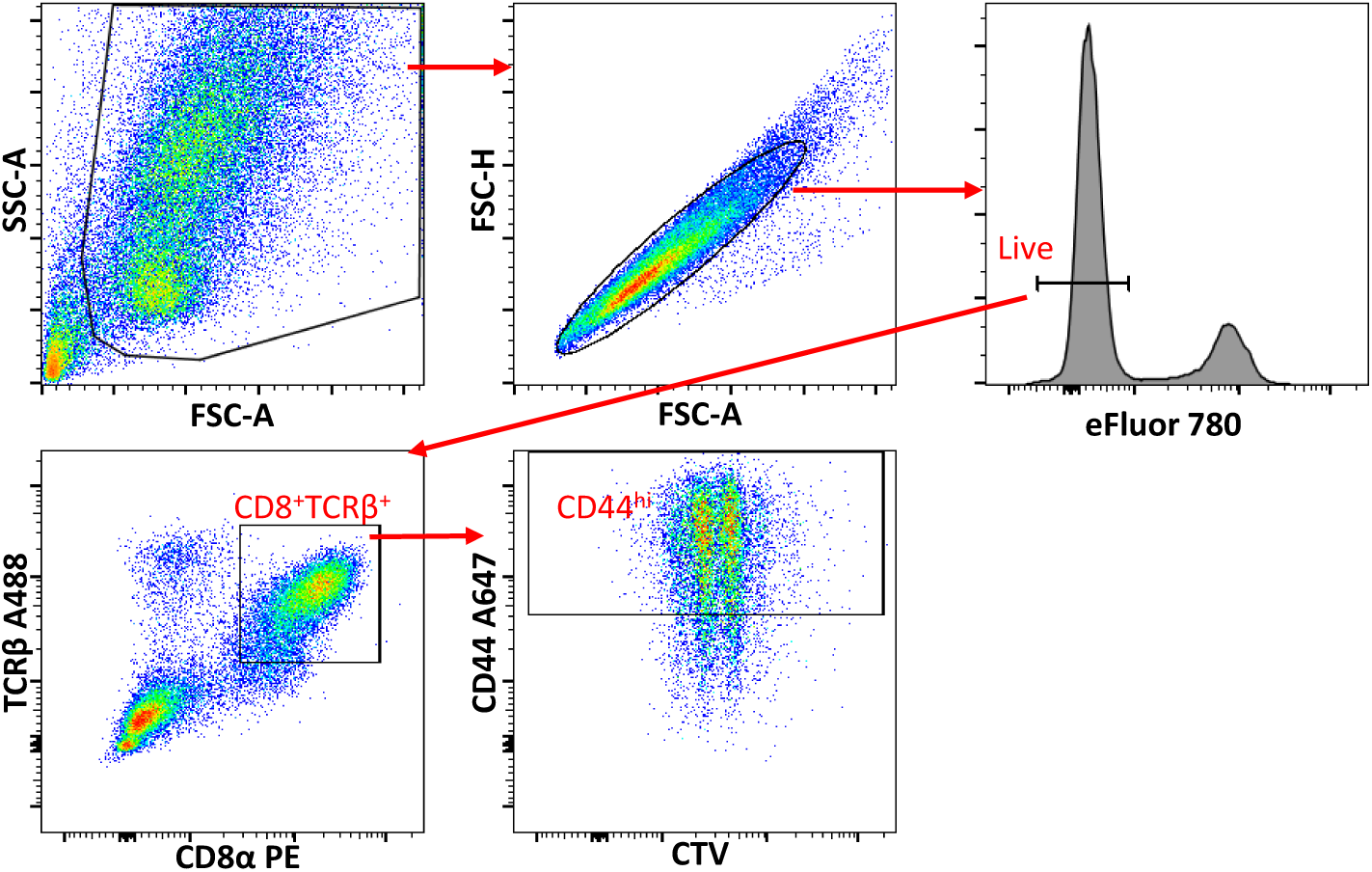
Flow cytometry gating strategy for CD8^+^ T cells *ex vivo*. Any unstable portions of the run were gated out prior to analysis. *Ex vivo* cultured CD8^+^ T cells were gated first with scatter gating (FSC-A vs. SSC-A) to exclude debris followed by single cell gating (FSC-A vs. FSC-H) to remove doublets/clumps. For proliferation assay, CD8^+^ T cells were identified as CD8^+^TCRβ^+^ cells within live population. CTV proliferation dye dilution was further characterized in CD44^hi^ population within these cells.

**Figure S11.**
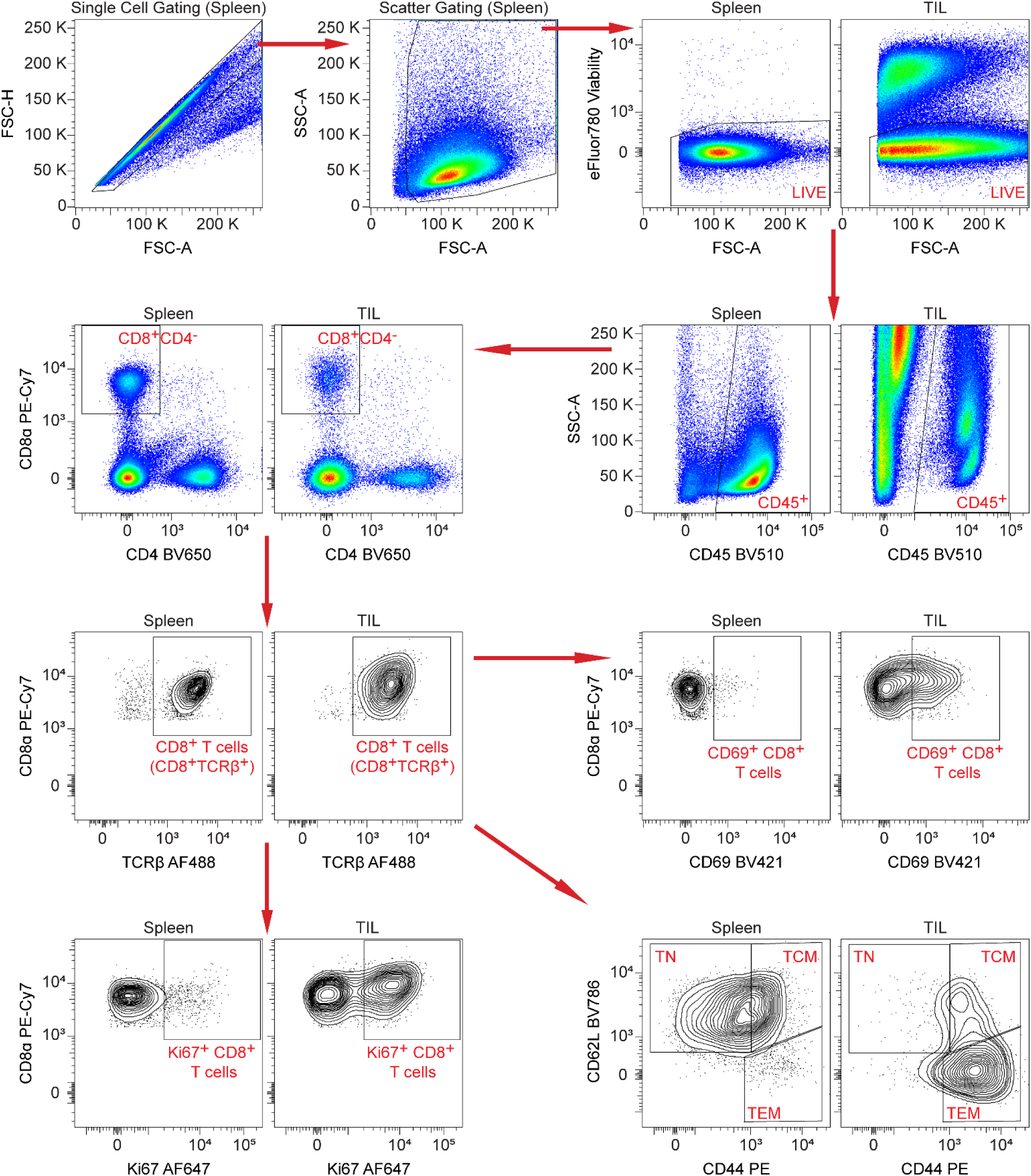
Flow cytometry gating strategy for CD8^+^ T cell profiling in spleen, TIL, and DLN. Any unstable portions of the run were gated out prior to analysis. After single cell gating (FSC-A vs. FSC-H) to remove doublets/clumps and scatter gating (FSC-A vs. SSC-A) to exclude debris, CD8^+^ T cells were identified as CD8^+^CD4^-^TCRβ^+^ cells within the live, CD45^+^ immune cell population. CD8^+^ T cells were further profiled to identify proliferating (Ki67^+^), newly activated (CD69^+^), naïve (TN, CD62L^lo^CD44^lo^), central memory (TCM, CD62L^hi^CD44^hi^), and effector/effector memory (CD62L^lo^CD44^hi^) CD8^+^ T cells. Example plots are shown for spleen and TIL for all gating following single cell and scatter gating.

